# A dedicated motif in human polymerase gamma enables DNA synthesis through replication roadblocks

**DOI:** 10.64898/2026.09.27.754746

**Authors:** Samuel Miguez-Amil, Araceli Grande-García, Plaza G.A. Ismael, María Moreno-Morcillo, Ester Casajús-Pelegay, Emma Arean-Ulloa, Carlos Chacón-Sánchez, Grzegorz L. Ciesielski, Borja Ibarra, Rafael Fernandez-Leiro

**Affiliations:** Spanish National Cancer Research Centre (CNIO), Melchor Fernández Almagro 3, 28029 Madrid, Spain; IMDEA Nanoscience, Cantoblanco, 28049 Madrid, Spain; Department of Biology, University of North Florida, Jacksonville, FL, United States

**Keywords:** mitochondria, mtDNA, DNA polymerase gamma, Template Stabilising Motif (TSM), strand displacement, RNA-DNA hybrids, G-quadruplexes, cryo-EM, single-molecule analysis

## Abstract

Mitochondrial DNA (mtDNA) maintenance is essential for cellular homeostasis, and defects in mtDNA replication are linked to a broad spectrum of mitochondrial diseases. During replication, DNA polymerase γ (Polγ) must traverse duplex junctions and stable secondary structures, yet how the human enzyme overcomes these barriers remains incompletely understood. Here, cryo-electron microscopy structures of Polγ bound to forked DNA, G-quadruplex (G4)-containing DNA and DNA bound to mitochondrial single-stranded DNA-binding protein (mtSSB) reveal a common template-entry path along the catcher domain across distinct substrate and active-site configurations. Within this domain, an arginine-rich helix containing R1026, R1030 and R1034 constitutes a Template Stabilising Motif (TSM). Biochemical reconstitution and DNA-binding experiments show that disrupting the TSM selectively impairs strand displacement, RNA-DNA hybrid displacement and synthesis through G4-forming sequences, while largely preserving synthesis on unstructured templates. Single-molecule optical-tweezers experiments further show that mechanical destabilisation of the fork partially restores mutant strand-displacement activity, whereas force or mtSSB restores primer-extension kinetics on ssDNA templates. Together, these findings establish the role of the TSM in maintaining productive template engagement and identify template stabilisation as a common mechanism enabling Polγ to traverse structurally diverse barriers in mtDNA.

## 1. INTRODUCTION

Mitochondrial diseases are a significant public health concern, with 1 in 200 individuals carrying pathogenic mtDNA mutations^1–3^. These conditions span a broad clinical spectrum, ranging from late-onset local muscle deficiencies (e.g., progressive external ophthalmoplegia) to severe, early-onset compound multisystemic disorders (e.g., Alpers syndrome). Moreover, mtDNA maintenance defects have also been associated with neurodegenerative diseases, metabolic syndromes, and cancer^4–6^.

The key elements responsible for maintaining mtDNA integrity are the components of the mtDNA replication machinery, known as the replisome, which includes the DNA polymerase gamma holoenzyme (Polγ), comprising the catalytic subunit (PolγA) and two copies of the accessory processivity factor (PolγB), the Twinkle helicase, and the mitochondrial single-stranded DNA-binding protein (mtSSB)^7^. The coordinated action of these factors facilitates processive DNA synthesis *in vitro*^8^ and is essential for mtDNA maintenance *in vivo*^9–11^. Although key steps of the mtDNA replication process are not fully understood, prevailing models support an asynchronous, strand-displacement process in which the synthesis of the nascent heavy strand occurs concomitantly with the displacement of the parental heavy strand^7,12^. During this process, the exposed strand is thought to be organised by either mtSSB molecules^13,14^ or RNA fragments incorporated through a bootlace-like mechanism^15,16^. As replication progresses, continued displacement of the parental heavy strand leads to exposure of the origin of light-strand replication (oriL), which adopts a stem-loop structure that triggers a secondary initiation event. Light-strand synthesis then proceeds in the opposite direction, mediated by Polγ acting on the previously displaced and protected strand as template, and involves replication across templates that may remain coated by mtSSB or contain RNA-DNA hybrids and stable secondary structures. Replication continues until the converging forks meet, followed by primer removal and termination of DNA synthesis. The correct assembly and functioning of the replisome are essential for mtDNA maintenance, and mutations in nuclear genes encoding replisome factors, particularly *POLG*^4^, which encodes PolγA, are the primary contributors to mtDNA replication-related pathologies^4^.

As the primary polymerase for mtDNA replication, Polγ maintains genome fidelity by coupling synthesis with proofreading, helping prevent the accumulation of mutations^17^. In addition to its canonical polymerase and exonuclease functions, Polγ has been shown to displace mtSSB from the template strand^18^, perform strand-displacement DNA synthesis^19,20^, bypass complex DNA structures^21^, and conduct translesion DNA synthesis^22,23^. Strand displacement activity is particularly critical at multiple stages of mtDNA replication, such as D-loop formation^12^, removal of RNA primers in coordination with RNase H1 and other enzymes^24–26^, and synthesis of the light strand following OriL exposure^12^. In models requiring RNA incorporation along the displaced strand, Polγ must also negotiate RNA-DNA hybrid intermediates during replication. Apart from these, Polγ faces other naturally occurring DNA roadblocks encountered during mitochondrial genome replication, such as G-quadruplexes (G4)^27^ on the template strand. G4-forming sequences are abundant within human mtDNA, particularly, the HSP1 region and 10220-10169 sequence can adopt stable G4 conformations *in vitro*^21,28,29^ and have been shown to form *in vivo*^29–31^. Moreover, Polγ has been reported to display variable tolerance toward different mitochondrial G4s^20^.

Studies of the homologous T7 replisome have suggested that the replicative polymerase provides the main driving force for DNA unwinding^32^. A similar mechanism may operate in the human mitochondrial replisome, where the activity of Polγ has been shown to be essential for promoting DNA unwinding by Twinkle, a helicase with intrinsically weak and strongly autoregulated activity^33^. Collectively, Polγ activities are critical for the proper function of the mtDNA replisome and for preventing DNA breaks and deletions^34^, thereby maintaining the integrity of mtDNA and mitochondria^34^.

Despite extensive structural and biochemical work on Polγ^17,35–42^, we still lack a mechanistic explanation for how the human enzyme overcomes duplex junctions or stable secondary structures during mtDNA replication. Here, we use cryo-EM to capture Polγ in multiple barrier-engaged states, including strand-displacement DNA synthesis, interaction with a G4 DNA substrate, and association with mtSSB (Fig. 1). By integrating these structural snapshots with *in vitro* functional studies and single-molecule analysis, we define the function of an arginine-rich helix within the previously described catcher domain that is largely dispensable for synthesis on unstructured templates but becomes critical when Polγ must maintain productive engagement with transiently unpaired template DNA during displacement of DNA duplexes and RNA-DNA hybrids and replication through G4-forming sequences. Together, these findings establish template stabilisation as a general mechanism by which Polγ traverses structurally diverse barriers during mtDNA replication and provide a framework for interpreting disease-linked variants that affect this interface.

**Figure 1.**
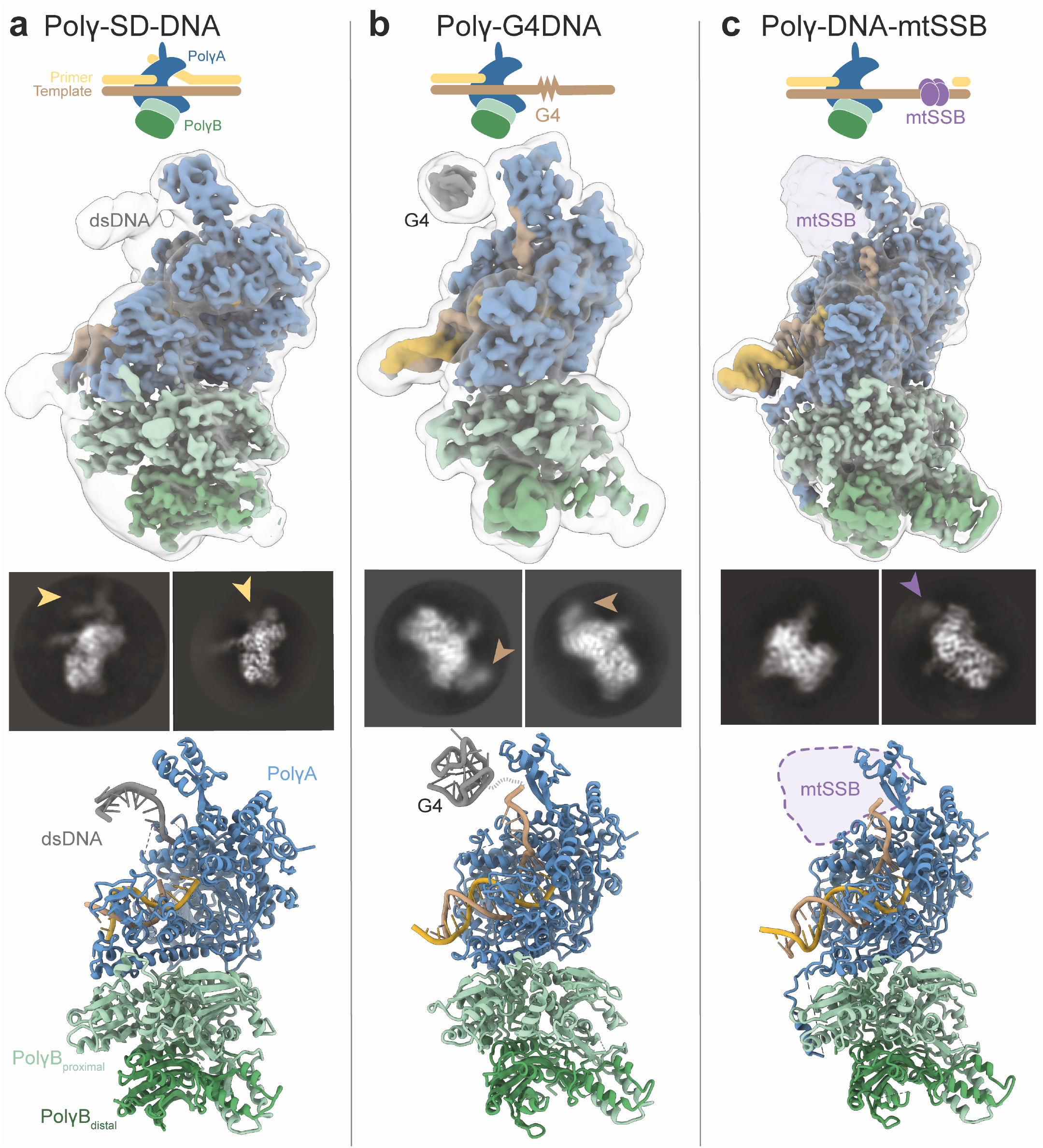
Cryo-EM structures of Polγ bound to distinct replication substrates. **a**, Cryo-EM reconstruction of the Polγ holoenzyme bound to a strand-displacement DNA substrate (Polγ-SD-open). A schematic of the substrate is shown above, representative 2D class averages are shown below the reconstruction, and the corresponding atomic model is shown at the bottom. Additional density compatible with the incoming duplex DNA is indicated in the 2D classes. **b**, Cryo-EM reconstruction of Polγ bound to a G4-containing DNA substrate. The corresponding substrate schematic, representative 2D class averages and atomic model are shown. Density corresponding to the G4 is indicated. **c**, Cryo-EM reconstruction of Polγ bound to DNA in the presence of mtSSB. The corresponding substrate schematic, representative 2D class averages and atomic model are shown. Additional density compatible with mtSSB is indicated in the 2D classes and by a dashed outline in the atomic model. PolγA is shown in blue, the two PolγB subunits in green, and DNA in yellow/brown. Regions shown in grey were not included in the final atomic models because the local density was insufficient for unambiguous model building; reference models were nevertheless fitted into the density and are shown to indicate their approximate positions.

## 2. RESULTS

### 2.1. Cryo-EM snapshots capture Polγ in multiple replication states

To understand how human Polγ progresses through DNA forks and other barriers encountered during mitochondrial DNA replication, we determined cryo-EM structures of the Polγ holoenzyme bound to a primed forked DNA substrate, a G4-containing primed template, and a long ssDNA template coated by mtSSB (Fig. 1, Supplementary Table 1).

We determined the structure of Polγ bound to the forked DNA substrate in the presence of dNTPs using a substrate that allowed the polymerase to extend the primer and encounter the duplex junction without artificial stalling or chemical trapping. This design enabled Polγ to sample barrier-engaged states under active turnover conditions. The dataset exhibited substantial compositional and conformational heterogeneity. Initial 3D classification identified a population of particles lacking DNA, from which we determined the structure of apo Polγ (Polγ-apo) at 3.2 Å resolution (Supplementary Fig. 1). Further classification of the DNA-bound particles yielded a reconstruction at 3.1 Å resolution, comprising approximately 20% of the DNA-bound particles, in which the primer terminus occupied an intermediate position resembling previously described configurations^17,40^. We refer to this reconstruction as Polγ-SD-open. A second class, comprising the remaining approximately 80%, yielded a reconstruction at 3.7 Å resolution in which the protein was sufficiently resolved for refinement, but the primer-terminal DNA density did not allow for confident assignment to a polymerase, exonuclease or intermediate configuration (Polγ-SD-close, Fig. 1a and Supplementary Fig. 1). The poorly resolved DNA in this class is consistent with heterogeneous substrate positioning at the primer-template junction.

To capture Polγ engaging a G4-containing DNA, we incubated the polymerase with a primer-template substrate harbouring a stable G4-forming sequence derived from the mitochondrial non-coding region (Supplementary Table 2) in the presence of dNTPs, substituting dATP with ddATP to block extension right before the G4 structure. This dataset yielded multiple structures with different registers of the DNA substrate with resolutions ranging from 3.2 to 3.4 Å (Fig. 1b and Supplementary Fig. 2). Finally, using a gapped primer-template DNA substrate with an internal 60-nt-long ssDNA section (Supplementary Table 2), we obtained a 2.4 Å reconstruction of Polγ bound to this substrate in the presence of mtSSB, with additional density compatible with mtSSB bound along the template strand (Fig. 1c and Supplementary Fig. 3).

The cryo-EM maps show well-defined density for the PolγA and PolγB components in all reconstructions, although the PolγB dimer exhibits greater flexibility (Supplementary Figs. 1-3). The dominant motion is centred on the PolγA thumb domain (residues 441-476 and 785-815), which acts as a hinge relative to the PolγB dimer. This motion is markedly reduced upon DNA binding, resulting in more homogeneous particle populations and improved resolution in the DNA-bound reconstructions. Masked refinements focused on PolγA and PolγB further improved the local quality of the corresponding maps (Methods and Supplementary Figs. 1-3). Density for the DNA regions within the catalytic cleft was generally well resolved, whereas more distal elements, including the incoming duplex, the G4 and mtSSB, displayed greater conformational heterogeneity and lower local resolution. These densities supported their positioning relative to Polγ and, in the case of the G4, were compatible with a folded quadruplex, but did not support de novo atomic modelling. We fitted reference models into these regions to illustrate their approximate positions, but did not subject them to atomic refinement.

The overall architecture of the holoenzyme in all reconstructions closely resembles previously reported Polγ structures^17,35–38,40^. Among the reconstructions with confidently resolved primer DNA ends, the primer terminus occupies an intermediate position in the Polγ-SD structure and the polymerase active site in the G4-containing and DNA-mtSSB-bound complexes. In the strand-displacement reconstruction, the intermediate configuration is accompanied by the previously described conformational rearrangements of the catalytic subunit associated with repositioning the primer terminus between catalytic states^17,40^. In the second strand-displacement class, the primer-terminal DNA remains insufficiently resolved for assignment to a specific catalytic configuration; however, the configuration of the DNA as it exits the active site partially resembles previously reported Polγ structures in the exonuclease state^17,40^.

The enrichment of intermediate configuration in the SD structure, rather than the polymerase configuration, is consistent with incoming duplex shifting the conformational equilibrium of Polγ away from productive synthesis. This interpretation is consistent with single-molecule studies showing that fork-regression pressure associated with DNA reannealing at the junction promotes polymerase pausing and stalling^19^, and with bulk biochemical studies linking these behaviours to idling and exonuclease processing^43,44^. By contrast, the DNA occupies the polymerase active site in the G4-containing and mtSSB-bound reconstructions, consistent with the absence of regression pressure from an incoming DNA duplex in these substrates (Fig. 1b,c).

Despite these rearrangements, the resolved portions of the incoming template converge on a common entry path adjacent to the previously described catcher domain^42^. In the strand-displacement dataset, the initial 2D class averages showed additional density extending from this region towards the template strand at the duplex junction (Fig. 1a), prompting the detailed analysis of the template-engagement interface described below.

### 2.2. A structural element in Polγ engages the incoming DNA template

A flexible helix-loop-helix element of PolγA (residues 991-1051) projects from the fingers domain (Fig. 2a,b). Initially proposed as a specialised extension of the Polγ fingers domain^45^, this region is now termed the catcher domain^42^, following the description of a structurally analogous element in the yeast mitochondrial polymerase Mip1^41^. In our reconstructions, the resolved segments of the incoming template follow a common entry path along this domain despite differences in substrate and primer positioning, encompassing intermediate state in the strand-displacement dataset and polymerase states in the G4-containing and mtSSB-bound datasets. Within the catcher domain, we focus on the arginine-rich helix containing R1026, R1030 and R1034, which we term the template stabilising motif (TSM) based on the functional evidence presented below.

**Figure 2.**
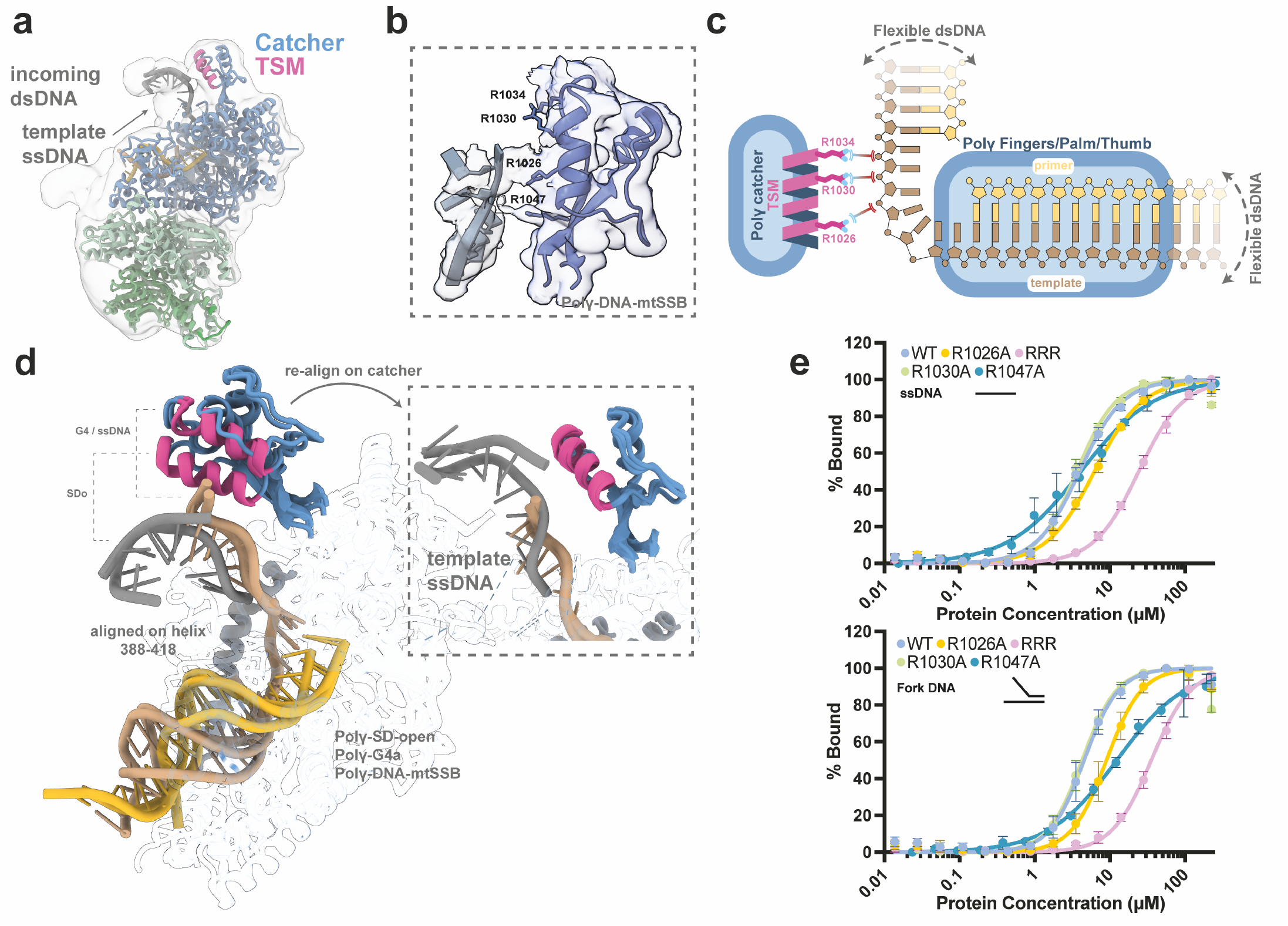
The Polγ catcher domain engages the incoming template through a template stabilising motif. **a**, Overview of Polγ bound to a strand-displacement DNA substrate, showing the incoming duplex and template ssDNA. PolγA is shown in blue, PolγB in green and the template stabilising motif (TSM) in magenta. The incoming duplex is represented by a reference model fitted into low-resolution density to illustrate its approximate position. **b**, Close-up of the catcher domain and adjacent template DNA with the corresponding cryo-EM density. Residues R1026, R1030 and R1034 within the TSM, together with R1047, are indicated. **c**, Schematic illustrating the trajectory of the incoming template along the TSM and into the catalytic core. The template and primer strands are shown in brown and yellow, respectively. Dashed arrows indicate flexibility of the distal duplex regions. **d**, Superposition of Polγ-SD-open, Polγ-G4a and Polγ-DNA-mtSSB, aligned on PolγA residues 388-418, a helix within the fingers domain that doesn’t contact DNA at any point. The catcher domain and TSM are highlighted in blue and magenta, respectively. The inset shows the same structures realigned on the catcher domain, illustrating a similar local template trajectory despite differences in catcher positioning relative to the polymerase core. **e**, Fluorescence-polarisation analysis of DNA binding by isolated catcher-domain constructs (PolγA residues 991-1051), comparing WT, R1026A, R1030A, R1047A and the R1026A/R1030A/R1034A triple mutant (RRR). Binding to ssDNA and forked DNA is shown in the upper and lower plots, respectively. Apparent K_D_ values for ssDNA and forked DNA, respectively, were 4.59 ± 0.98 and 3.95 ± 0.45 μM for WT; 9.12 ± 1.94 and 6.19 ± 0.46 μM for R1026A; 35.72 ± 4.00 and 24.28 ± 2.43 μM for RRR; 4.30 ± 0.68 and 3.64 ± 0.40 μM for R1030A; and 12.53 ± 2.34 and 4.17 ± 0.92 μM for R1047A. Values represent means from six independent experiments for WT, R1026A, R1030A and RRR, and three for R1047A; error bars indicate SD. Lines represent fitted binding curves.

Residues R1026, R1030, R1034, and R1047 create a surface close to the ssDNA template in all structures, which stretches out from the catalytic site parallel to this motif, suggesting potential interactions between this motif and the DNA (Fig. 2b-d). Indeed, we observe density consistent with contacts between side chains in helix 1022-1034 and the DNA phosphate backbone, as well as a further contact involving R1047 closer to the fingers domain (Fig. 2b-d). To test whether this motif was able to bind DNA and the specific contribution of each residue, we cloned and purified variants R1026A, R1030A, R1034A, and R1047A, along with a triple-arginine mutant R1026A-R1030A-R1034A (hereafter referred to as RRR) and a Polγ construct lacking the whole motif (Polγ-Δ991-1051). We purified these variants as isolated motifs (MBP-catcher; see methods for details) to test the specific contribution of the motif, and as reconstituted holoenzyme variants with the full-length versions of PolγA carrying these mutations to test their impact on the holoenzyme DNA-binding capacity (Supplementary Fig. 4a-c). Notably, all full-length variants retain thermal stability similar to that of the WT protein (Supplementary Fig. 4d), indicating that the mutations do not cause a detectable change in overall thermal stability. To corroborate this in the context of the short constructs, we conducted 50 ns molecular dynamics simulations on the motifs with the different mutations, which suggest similar stability across all variants (Supplementary Fig. 4d, 5).

Subsequently, we assessed DNA-binding by fluorescence polarisation using FAM-labelled DNA substrates (Supplementary Table 2, Fig. 2e). The isolated catcher-domain construct bound ssDNA and forked DNA with apparent K_D_ values of 4.59 and 3.95 μM, respectively. Individual substitutions had modest or substrate-dependent effects: R1030A retained apparent affinities similar to WT, R1026A showed approximately twofold and 1.6-fold increases in apparent K_D_ for ssDNA and forked DNA, respectively, and R1047A showed an approximately threefold increase for ssDNA but little change for forked DNA. In contrast, RRR exhibited approximately eightfold and sixfold increases in apparent K_D_ for ssDNA and forked DNA, respectively, supporting a contribution of the TSM basic surface to DNA binding. Mutations in the full-length proteins also reduced apparent DNA-binding affinity, particularly in the triple mutant and catcher-domain deletion construct (Supplementary Fig. 4g), indicating that this region contributes to DNA binding in the intact polymerase.

These results indicate that the positively charged residues R1026, R1030 and R1034 within the TSM contribute directly to DNA binding, consistent with their position along the template-facing surface in the cryo-EM structures. Across the different reconstructions, the resolved template segments adopt a similar sharply bent trajectory as they pass the TSM and approach the catalytic core (Fig. 2d). Notably, in the strand-displacement reconstructions, between approximately 5 and 8 nucleotides of the template remain unpaired between the duplex junction and the primer terminus, despite the reannealing pressure exerted by the incoming duplex. Beyond this segment, the density becomes diffuse, consistent with increased DNA flexibility. Nevertheless, refinement centred on this region recovered low-resolution density compatible with the incoming duplex ahead of the fork (Fig. 2a). Together, these observations place the TSM along the exposed template segment connecting the duplex junction to the catalytic core.

### 2.3. Strand-displacement activity depends on Polγ’s template DNA binding motif

To define the contribution of the catcher domain to strand displacement, we generated a panel of Polγ variants carrying substitutions within and outside the TSM, together with a catcher-domain deletion construct (Polγ-Δ991-1051), and assessed their primer-extension (PE) and strand-displacement (SD) activities (Fig. 3a,b and Supplementary Fig. 4f,h).

**Figure 3.**
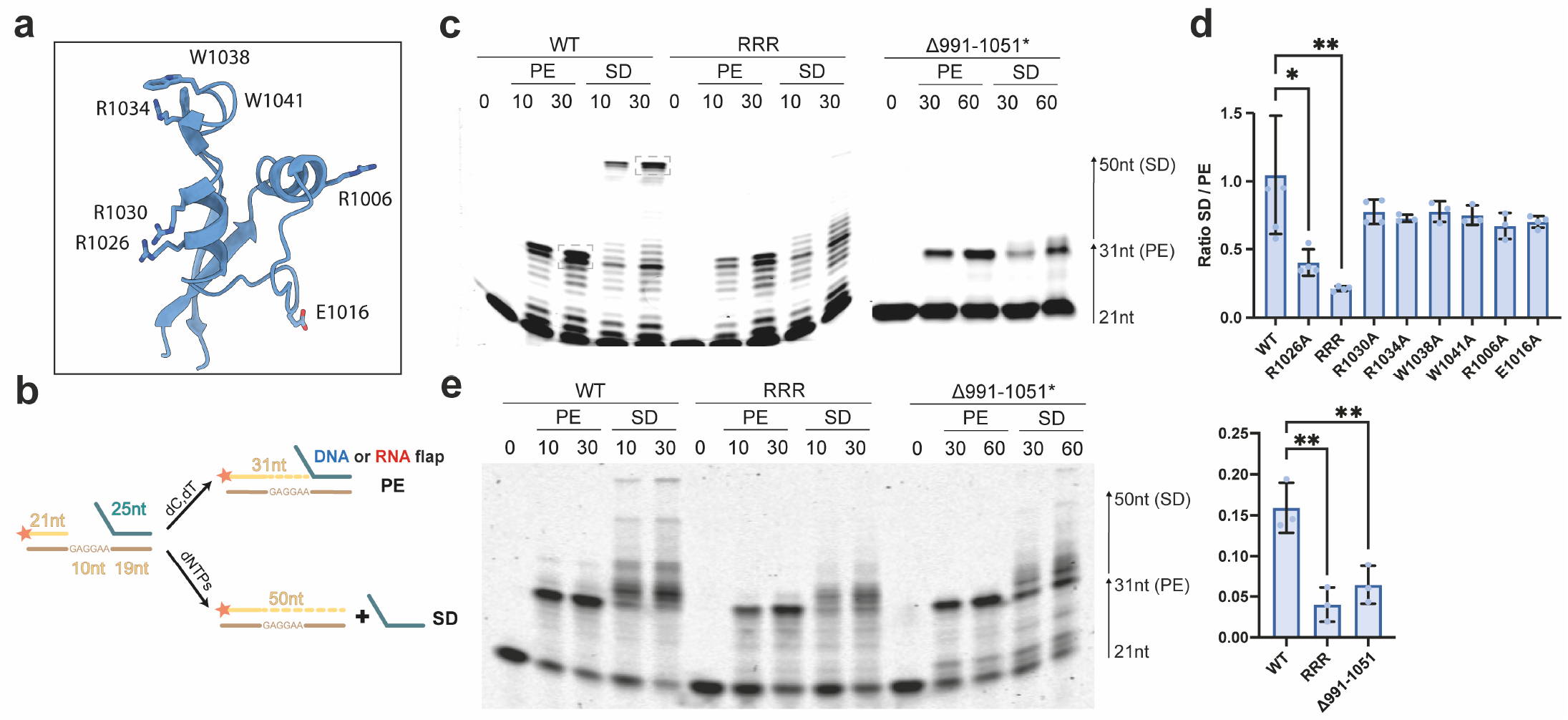
The template-engaging element is required for efficient strand displacement. **a**, Structural representation of the template-engaging element highlighting residues analysed by mutagenesis. **b**, Schematic representation of the DNA substrates and nucleotide conditions used to distinguish primer-extension (PE) and strand-displacement (SD) activities. **c**, Representative PE and SD assays performed with WT Polγ, the RRR mutant and the Polγ-Δ991-1051 mutant using DNA-DNA hybrid substrates at the indicated reaction times. Left, products generated in the primer-extension and strand-displacement reactions, detected using IRDye700-labeled primers and analysed by electrophoresis on 15% TBE-urea polyacrylamide gels. Right, quantification of strand-displacement activity relative to primer-extension activity based on independent reactions analysed by capillary electrophoresis using FAM-labelled primers (n ≥ 3 biological replicates). Dashed boxes illustrate the regions used for quantification both in the gels and capillary electrophoresis. Asterisk indicates protein concentrations 6-fold higher than those used for the other proteins. **d**, Corresponding PE and SD assays using RNA-DNA hybrid substrates. The ratio between PE and SD is presented on the graphs as mean ± standard deviation from 3 biological independent experiments; individual data points represent independent experiments. Asterisk indicates protein concentrations 6-fold higher than those used for the other proteins. Statistical significance relative to WT was assessed using a two-tailed paired Student’s t-test (*P ≤ 0.05; **P ≤ 0.01).

WT Polγ efficiently performed both PE and SD activities (Fig. 3c). In contrast, Polγ-Δ991-1051 exhibited almost no detectable SD activity, whereas PE was only partially impaired. This reduction in PE may reflect both broader structural perturbations caused by deletion of the complete catcher domain and loss of its contribution to template organisation during primer extension.

Substitutions targeting the TSM impaired SD while leaving PE largely unaffected, whereas substitutions elsewhere in the catcher domain had much milder effects on SD (Fig. 3c,d and Supplementary Fig. 4f). The SD/PE ratios of R1026A, RRR and Polγ-Δ991-1051 were significantly lower than those of WT. In particular, the R1026A and RRR variants exhibited markedly impaired SD activity while retaining PE activity comparable to WT, corresponding to reductions of 62% and 80% in the SD/PE ratio, respectively. Together, these results support a role for the catcher domain in strand-displacement synthesis and identify the TSM as the principal contributor to this activity within the domain.

We next asked whether the TSM also contributes to strand displacement across RNA-DNA hybrids, which arise during mitochondrial replication through incorporation of RNA fragments on the displaced strand and during primer removal. As shown in Figure 3e, WT Polγ can efficiently process RNA-DNA hybrids. In contrast, the RRR and Polγ-Δ991-1051 mutants show reduced activity across RNA-DNA hybrids (78% and 62% reduction in SD/PE ratio, respectively), indicating that the requirement for TSM extends to strand displacement across RNA-DNA hybrids.

### 2.4. Mechanical destabilisation of base pairing ahead of the polymerase rescues strand-displacement activity

We next used optical tweezers to characterise the effect of mutations that disrupt TSM function (RRR variant) on the SD and PE real-time kinetics of Polγ. We conducted these experiments as a function of mechanical force applied to the DNA, which has been shown to decrease base-pair stability and promote the activities of Polγ and other polymerases^18,19,46^. Briefly, a primer-template DNA molecule (for PE) or a forked DNA structure (for SD) was tethered between functionalised micrometre-sized polystyrene beads. Under constant mechanical force, as the polymerase unwinds the fork during SD or converts ssDNA to dsDNA during PE, the bead-to-bead distance changes (Fig. 4). These changes in tether extension were converted into the number of nucleotides unwound or incorporated using the corresponding extension per nucleotide at each force (see methods for more details). The WT polymerase presented a robust SD activity above 5 pN of force applied to the opposite strands of the DNA fork (Fig. 4a). Under these conditions, the average processivity (number of replicated nucleotides), average velocity and time in pause state per nucleotide at low (6 pN) and high (10 pN) forces were compatible with those measured previously for this polymerase^19^. In contrast, the RRR variant showed no detectable activity at forces below 9 pN. Activity became detectable only at 10 pN, a force that reduces the energetic barrier to fork opening but is insufficient to unfold the hairpin in the absence of polymerase^47^. Mechanical destabilisation therefore partially bypassed the defect caused by TSM disruption. However, even under these conditions, the RRR variant exhibited altered SD kinetics, showing reduced processivity and velocity, and increased time per nucleotide in the pause state, relative to WT polymerase (Fig. 4a). These results can be interpreted in light of previous single-molecule experiments showing that mechanical force applied to the complementary DNA strands suppresses fork regression, thereby decreasing competition with Polγ for engagement of the template strand^19^. Stable engagement of the template strand by WT Polγ therefore supports strand-displacement activity even at low force. In contrast, the reduced template affinity of the RRR mutant is sufficient to sustain strand displacement only when applied force decreases base pair stability and counteracts fork regression.

**Figure 4.**
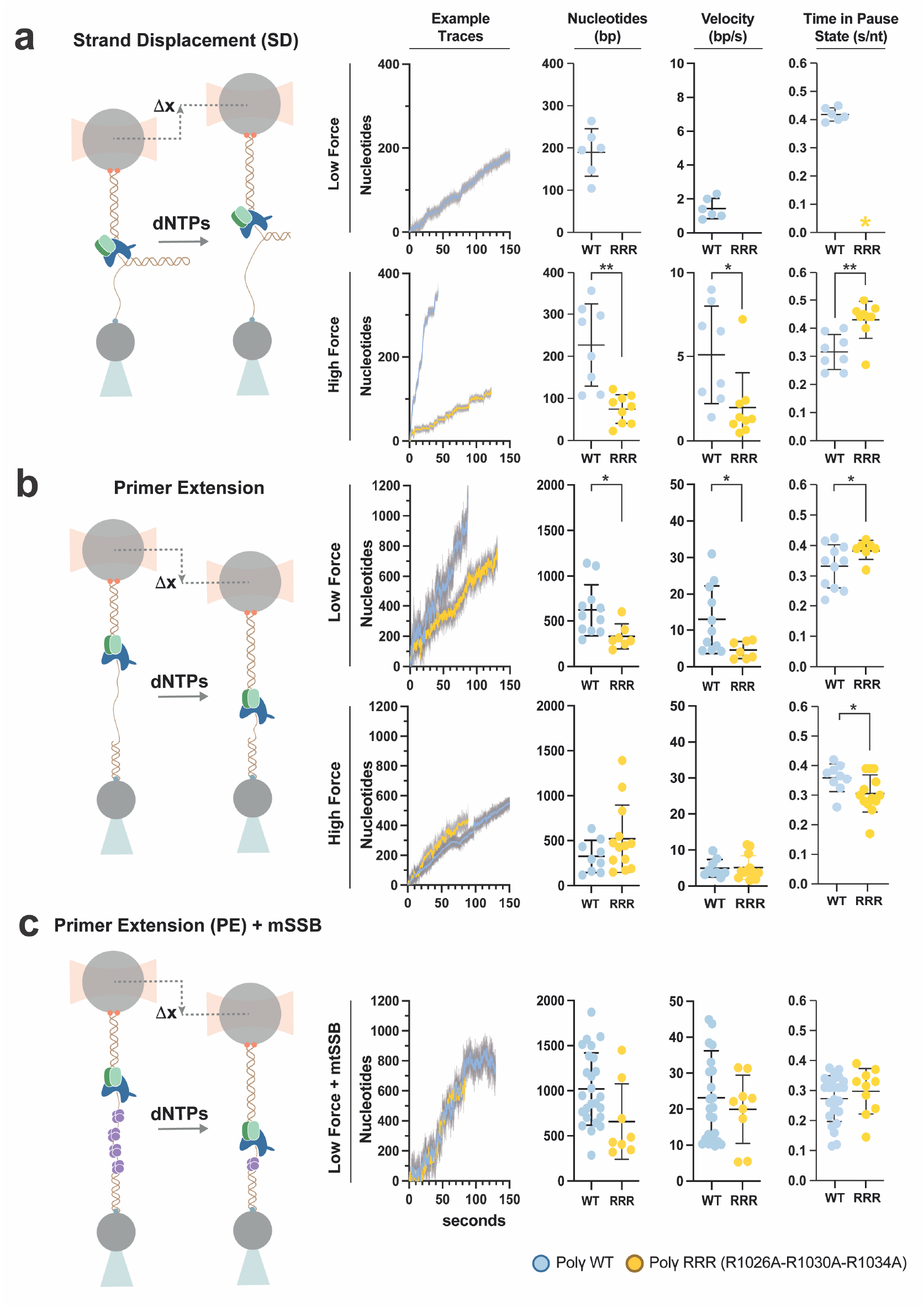
Mechanical destabilisation of DNA rescues TSM-defective Polγ. **a**, Single-molecule optical-tweezers analysis of strand-displacement activity. The experimental configuration is shown on the left. The four panels on the right show representative traces for WT and RRR Polγ at low and high force, and quantification of processivity, velocity, and time spent in the pause-state duration per nucleotide under the indicated force conditions. **b**, Single-molecule optical-tweezers analysis of primer-extension activity by WT and RRR Polγ. Representative traces and corresponding measurements of processivity, velocity and pause-state duration are shown at low and high force. **c**, Single-molecule measurements of primer-extension activity in the presence of mtSSB. Representative traces and corresponding measurements of processivity, velocity and pause-state duration are shown for WT and RRR Polγ. In all panels, individual measurements correspond to single-molecule trajectories. WT Polγ is shown in blue and RRR Polγ in yellow.

Analysis of primer-extension kinetics (Fig. 4b) showed that at low force (2 pN), the RRR mutant exhibited significantly reduced processivity and velocity, along with an increased pause-state duration per nucleotide, relative to WT Polγ. These defects were rescued to WT-like levels when force was applied to the DNA template (10 pN). Notably, primer-extension kinetics at low force were also restored by the addition of mtSSB (20 nM) (Fig. 4c). Because both mechanical stretching of the DNA and mtSSB binding destabilise secondary structures in the template ssDNA^18^, these findings further support the idea that stable DNA secondary structures ahead of the polymerase impair the activity of the RRR variant.

Together, these results demonstrate that the TSM promotes productive engagement of the template strand regardless of whether the energetic barrier arises from an incoming duplex during strand displacement or from secondary structures formed within the template ssDNA. These findings support a model in which the TSM organises and stabilises the incoming template, thereby facilitating synthesis through structured DNA.

### 2.5. The TSM facilitates replication through G4-forming template DNA

Prompted by these results, we next asked whether the TSM contributes to DNA synthesis through stable DNA secondary structures encountered during mitochondrial genome replication, such as G-quadruplex (G4)^27^ forming sequences in the template strand. We reconstituted primer-template substrates containing well-characterised mitochondrial G4-forming sequences, G4a and G4b, previously shown to impose barriers of different strengths to Polγ progression^21^ (Supplementary Table 2). DNA synthesis was compared between WT Polγ, the RRR variant and Polγ-Δ991-1051 (Fig. 5a).

**Figure 5.**
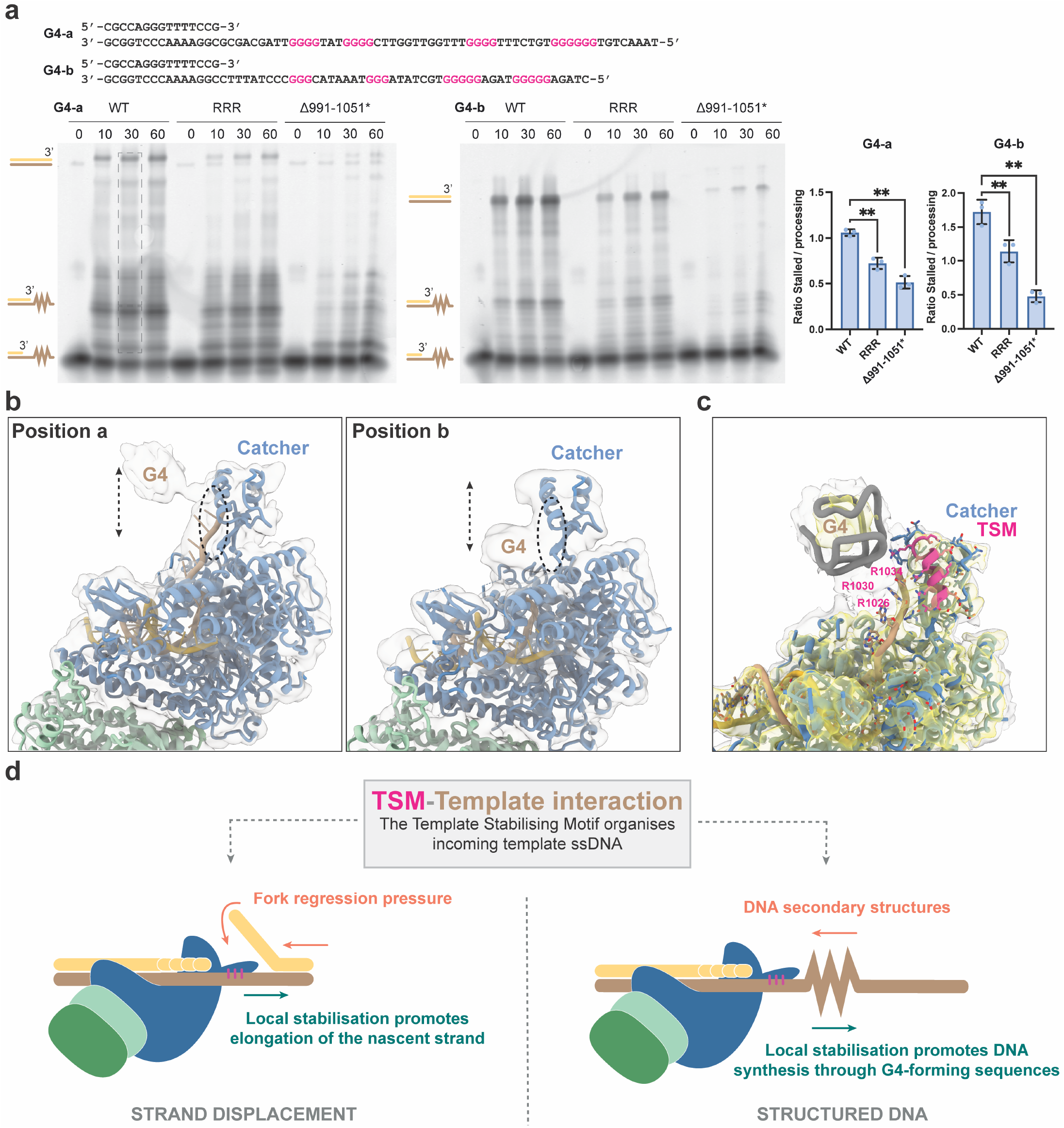
The TSM facilitates DNA synthesis through mitochondrial G4-forming sequences. **a**, Primer-extension assays using two mitochondrial G4-forming DNA substrates previously characterised as stronger (G4a) or weaker (G4b) barriers to Polγ progression. Reactions were performed with WT Polγ, RRR and Polγ-Δ991-1051 for the indicated times. Polγ-Δ991–1051 was assayed at a 6-fold higher protein concentration because of its reduced basal polymerase activity. G4 traversal was quantified as the ratio of fluorescence intensity from products extending beyond the G-rich sequence to that from products terminating before or within this region (dashed boxes). The graphs show the extension ratio between stalled and processed products at G4-forming sequences as mean ± SD from 3 independent biological experiments; individual data points represent independent experiments. Statistical significance relative to WT was assessed using a two-tailed paired Student’s t-test (*P ≤ 0.05; **P ≤ 0.01). Asterisk indicates protein concentrations 6-fold higher than those used for the other proteins. **b**, Cryo-EM structures of Polγ bound to G4-containing DNA, showing two representative positions of the G4 upstream of the polymerase active site. The TSM and G4 are indicated. **c**, Close-up view of the G4-containing template, showing the G4 upstream of the active site and the intervening ssDNA engaged by the TSM. **d**, Model illustrating the general mechanism by which the TSM organises the incoming template during strand displacement at duplex DNA and during synthesis through G4-forming sequences. Statistical significance relative to WT was assessed using a two-tailed paired Student’s t-test (*P ≤ 0.05; **P ≤ 0.01).

Because Polγ-Δ991-1051 showed reduced overall polymerase activity, it was assayed at a sixfold higher concentration to obtain products within a quantifiable range (Methods and Fig. 5a). G4 traversal was evaluated using the ratio between products extending beyond the G-rich sequence and products terminating before or within this region. For the more readily traversed G4b substrate, WT Polγ reached a processed-to-stalled product ratio of approximately 1.7, whereas both RRR and Polγ-Δ991-1051 showed lower ratios (1.1 and 0.5, respectively). The effect was more pronounced with G4a, for which both variants exhibited a disproportionate accumulation of stalled products relative to products extending beyond the G-rich sequence (Fig. 5a). Thus, the altered product distribution is not readily explained by reduced overall activity alone and supports a role for the TSM in promoting Polγ progression through G4-forming templates.

To rationalise these functional findings, we determined cryo-EM structures of Polγ bound to a primer-template containing a pre-folded G4 structure on the template strand (Fig. 1b, 5b and Supplementary Fig. 2). The folded G4 was positioned upstream of the polymerase active site, with an intervening ssDNA segment of approximately seven nucleotides following a path adjacent to the TSM. Multiple 3D classes captured alternative substrate configurations in which the G4 density occupied different positions relative to the active site, while the resolved intervening ssDNA retained a similar trajectory along the TSM surface (Fig. 5b,c). In the configuration with the G4 closest to the active site, the basic TSM residues were positioned close to what would be the phosphate backbone on the outer surface of the G4, suggesting that this interface may also accommodate the structured portion of the template. Together with the reduced G4 traversal exhibited by TSM-defective variants, these observations support a role for the TSM in maintaining template engagement as Polγ approaches the G4 barrier.

### 2.6. The catcher domain is not required for mtSSB displacement

MtSSB has been shown to stimulate Polγ primer extension activity, enabling the polymerase to achieve maximum velocity. Moreover, this effect is species-specific, as other ssDNA-binding proteins such as *E. coli* SSB do not stimulate Polγ to the same extent. This specificity implies a functional coupling between the polymerase and mtSSB. While previous work has suggested that this coupling is mediated by electrostatic repulsion^18,46,48^, the Polγ region responsible for this effect has not been identified.

In the Polγ-DNA-mtSSB reconstruction, Polγ and the DNA substrate were well resolved, whereas mtSSB remained diffuse, consistent with substantial positional heterogeneity and previous reports^45^. At lower map thresholds, additional density compatible with mtSSB was observed over the incoming ssDNA (Supplementary Fig. 6a). Negative-stain 2D classes of the same complex confirmed multiple orientations of mtSSB relative to Polγ, explaining its poor high-resolution definition in the cryo-EM reconstruction (Supplementary Fig. 6b, c).

Despite its high flexibility, we observed mtSSB density localising near the TSM, suggesting that this region may contribute to mtSSB interaction. To test this, we analysed primer extension by WT and Polγ-Δ991-1051 holoenzymes on a long ssDNA template in the absence or presence of mtSSB. Although Polγ-Δ991-1051 displayed lower overall activity, mtSSB did not specifically impair its activity relative to WT (Supplementary Fig. 6d). Consistent with this, optical tweezers experiments showed that mtSSB restored primer-extension kinetics of the TSM-defective RRR variant to WT-like levels rather than inhibiting its activity. Together, these results indicate that the TSM is not required for productive coupling with mtSSB during DNA synthesis.

## 3. DISCUSSION

Mutations in *POLG*, which encodes the catalytic subunit of Polγ, are associated with a wide spectrum of mitochondrial diseases, highlighting the central role of this enzyme in mtDNA replication and maintenance^4,7^. As the sole mitochondrial replicative DNA polymerase, Polγ must overcome several challenging DNA structures during mtDNA genome replication, including DNA fork structures, RNA-DNA hybrids, and G4-rich regions. Progression through these obstacles requires local remodelling of the incoming DNA to expose the template strand for synthesis. Recent studies have described catcher-DNA interactions in strand displacement by yeast Mip1 and the human Polγ catcher domain^41,42^; however, how the catcher-domain interface contributes to synthesis through different structural barriers remained incompletely understood. Here, we functionally define an arginine-rich helix within this domain as the template stabilising motif (TSM) and establish its contribution to synthesis through distinct structural barriers. Across our reconstructions, the resolved incoming template follows a common trajectory along the catcher-domain surface despite changes in substrate and active-site configuration. Together with the biochemical and single-molecule results, these structures support a model in which the TSM preserves productive template engagement during catalytic-state transitions and replication through structured DNA barriers, without requiring direct recognition of each barrier.

Functional analysis of multiple TSM variants showed that mutations affecting the positively charged residues that contact the template strand reduce DNA binding while largely preserving primer-extension activity. Single-molecule measurements independently showed reduced productive DNA engagement by the RRR mutant. Importantly, these mutants displayed pronounced defects in strand displacement, with substitutions at R1026, R1030 and R1034 (Supplementary Fig. 4) selectively impairing this activity and the triple RRR mutant exhibiting a severe strand-displacement defect despite near-WT primer-extension activity. Optical tweezers experiments further showed that the strand-displacement defect of the RRR mutant could be rescued by mechanical destabilisation of the fork. Likewise, impaired primer extension on structured ssDNA templates was restored either by force or by mtSSB, indicating that TSM becomes particularly important whenever structured DNA must be locally destabilised ahead of the polymerase.

Experiments using RNA-DNA hybrid substrates showed that TSM variants retain reduced strand displacement activity, consistent with the results obtained on duplex DNA. Unexpectedly, strand displacement was generally more efficient on RNA-DNA hybrid substrates than on duplex DNA forks for both WT Polγ and TSM mutants. Although the molecular basis for this difference remains unclear, it may reflect the distinct thermodynamic and structural properties of RNA-DNA hybrids compared with B-form duplex DNA^49,50^.

Our findings are consistent with previous studies showing that replicative holoenzymes typically exhibit rapid, processive primer extension but often inefficient strand-displacement DNA synthesis. This behaviour has been attributed to the regression force exerted by the fork against the polymerase’s forward motion, which promotes frequent sampling of exonuclease or idle states. Moreover, single-molecule studies have shown that stabilisation of the first unwound base pair and the stability of the fork ahead of the polymerase are the key determinants of strand-displacement DNA synthesis^19^. Importantly, the requirement for intrinsic strand-displacement capacity is not bypassed by the presence of a replicative helicase. Several stages of mtDNA replication proceed without Twinkle ahead of the polymerase, including RNA primer removal^23–25^, light-strand synthesis^12^, and D-loop formation^12^. Even at the heavy-strand fork, unwinding by Twinkle is strongly autoregulated and accelerated by DNA synthesis^31,32^, indicating that the polymerase itself contributes to duplex separation even when the helicase is present.

Notably, neutralisation of TSM charges (RRR) or deletion of the motif (Polγ-Δ991-1051) also selectively impaired replication across G4-forming sequences while largely preserving primer extension on unstructured templates. Furthermore, cryo-EM captured the G4 upstream of the polymerase active site, with the TSM engaging the incoming template ssDNA similarly to forked DNA substrates. Together with the biochemical data, these structures support a model in which the TSM organises and stabilises exposed template ssDNA during replication through G4-forming sequences. This model explains why disruption of the TSM or deletion of the complete catcher domain reduces progression beyond the G-rich region and links G4 traversal to the template-organising mechanism used during duplex strand displacement. Rather than recognising or unwinding each obstacle directly, the TSM may provide a common template-stabilising surface that helps Polγ remain productively engaged when DNA structure opposes polymerase progression.

In contrast to its central role in overcoming structured DNA substrates, the catcher domain is not required for functional coupling between Polγ and mtSSB. During mitochondrial DNA replication, mtSSB coats exposed ssDNA and stimulates Polγ primer extension, implying that the polymerase must efficiently displace or bypass mtSSB bound to the incoming template to sustain DNA synthesis^14,20,46,48,51^. Although cryo-EM analysis revealed that mtSSB localises close to the catcher, consistent with previous reports^45^, the interaction remained flexible and heterogeneous, precluding the identification of a stable protein-protein interface. Moreover, deletion of this domain did not cause additional mtSSB-dependent inhibition of DNA synthesis on mtSSB-coated templates, and optical tweezers experiments showed that mtSSB restored primer-extension kinetics of the RRR mutant to WT levels, indicating that it is dispensable for mtSSB displacement during DNA synthesis, and that mtSSB has a redundant role with the TSM as both help stabilise the ssDNA template ahead of the polymerase.

Among human DNA polymerases, a comparable insertion is found only in Polδ, which also performs strand displacement during Okazaki fragment processing^52^, suggesting a shared functional principle (Supplementary Fig. 7a). Similarly, the catcher domain of the yeast mitochondrial DNA polymerase Mip1^41^ occupies a similar position relative to the DNA-entry channel, although its local architecture differs from that of the corresponding region in human Polγ. More broadly, mitochondrial DNA polymerases from diverse eukaryotic lineages contain structurally distinct catcher/TSM-like insertions adjacent to the DNA-entry channel (Supplementary Fig. 7b, 8). Despite their structural diversity, these elements recurrently display a basic surface adjacent to the DNA entry channel, consistent with convergent structural solutions to the common challenge of organising incoming DNA for replication through structured templates. Consistent with this possibility, the closely related T7 DNA polymerase gp5 lacks an equivalent element and exhibits limited strand-displacement activity and strong exonuclease activity.

Given the enrichment of G-quadruplex-forming motifs within the mitochondrial control region and their association with replication pausing and deletion breakpoints^28,29^, together with the persistence of structured DNA intermediates during mtDNA replication, defects in TSM-mediated template stabilisation would be expected to increase fork stalling, promote idling and exonuclease cycling, and ultimately increase the risk of replication-associated deletions. Accordingly, *POLG* variants that perturb TSM charge, positioning or dynamics may disproportionately impair replication through structured DNA elements, including G4 motifs at HSP1 and other regulatory regions. Consistent with this view, a recurrent observation in mitochondrial disorders is that certain *POLG* mutations exhibit modest or substrate-specific effects *in vitro*, particularly in assays measuring primer extension on unstructured templates, yet result in severe cellular phenotypes. Such discrepancies suggest that these variants may compromise specialised functions required for replication under physiological constraints, including strand displacement and replication through structured DNA.

Notably, several disease-associated variants map directly within or adjacent to the catcher domain (residues 991-1051). Among these, substitutions at R1047 (R1047W/Q/L)^53,54^, associated with Alpers-Huttenlocher syndrome, progressive external ophthalmoplegia (PEO) and epilepsy, seem to result in accumulation of mtDNA deletions. Although the severe polymerase defect reported for R1047W^53,54^ has been attributed primarily to disruption of the palm domain and dNTP binding, we find that R1047 contacts the incoming template and that its substitution reduces DNA binding to a similar extent as other TSM substitutions, raising the possibility that impaired TSM-mediated template stabilisation may further contribute to the severity of these variants. However, the disease-associated substitutions are non-conservative and may perturb the element more severely than the alanine substitution. More generally, additional disease-associated variants within the catcher region (e.g., A1033V, V1044A, G1051R^3,55,56^), as well as variants structurally coupled to the catcher such as F907I^57,58^, may similarly perturb the positioning or dynamics of this motif.

Collectively, our findings support a unifying model in which the TSM organises incoming DNA into a synthesis-competent configuration by stabilising transiently unpaired template nucleotides ahead of the polymerase. Whether during strand displacement at replication forks or during replication through stable DNA secondary structures such as G-quadruplexes, this local stabilisation may favour DNA opening and productive nucleotide incorporation. (Fig. 5d). By facilitating progression through these challenging replication barriers, TSM-mediated template stabilisation may limit fork stalling and replication stress, thereby reducing the risk of replication-associated DNA breaks and deletions and contributing to mitochondrial genome integrity. More broadly, our findings suggest that local stabilisation of transiently unpaired template DNA by specialised structural elements may represent a general strategy by which replicative DNA polymerases overcome structured DNA barriers.

## 4. METHODS

### 4.1. Cloning & Site-directed mutagenesis

All molecular cloning and mutagenesis were performed following the *in vivo* DNA assembly method^59^. DNA sequences encoding PolγA(30-1239)-3xFlag and 10xHis-mtSSB(17-148)-StrepTag were synthesised as gBlocksTM (Integrated DNA Technologies, IDT). For PolγB expression, a codon-optimised *POLG2* Δ25 (25-485) insert was subcloned into the pETite N-His SUMO vector (Lucigen). PolγA was cloned into the pACEBac1^60^ vector for expression in insect cells, while mtSSB and the catcher of PolγA were cloned into the pRSFDuet-1 vector (Novagen) for bacterial expression. Refer to Supplementary table 3 for a detailed list of variants.

### 4.2. Protein expression

The recombinant baculovirus genomes containing the PolγA sequences were generated by transforming DH10EMBacY cells (Geneva Biotech) with pACEBac1. The purified bacmid was transfected into *Sf* 9 cells cultured in *ESF921* medium to produce recombinant baculoviruses for PolγA expression. Finally, 2 L of *Sf* 9 cells were infected with baculoviruses at a density of 3×10^6^ cells/ml and incubated at 27 °C for 3-5 days. The cells were then harvested and snap-frozen for subsequent purification. PolγB and mtSSB and the PolγA-catcher fused to MBP were expressed in *E. coli* BL21, Rosetta (DE3) and LOBSTR-BL21(DE3)-RIL cells, respectively, using 6 L of LB medium (2L for catcher-MBP). When the culture reached an OD_600_ of 0.6, protein expression was induced with 1 mM IPTG for 18 hours at 18 °C.

### 4.3. Protein purification

Cells expressing wild-type or mutant versions of PolγA were lysed using a loose pestle in buffer 1 (50 mM HEPES pH 7.5, 500 mM KCl, 10 mM MgCl_2_, 0.1 mM EDTA, 0.1% Triton X-100, 10% glycerol, 2 mM β-mercaptoethanol, and cOmplete™ Protease Inhibitor Cocktail (Roche)) and followed by sonication (5 minutes; 3 s ON/6 s OFF cycles at 37% amplitude). The clarified supernatant (50,000 x *g* for 45 min) was incubated with anti-FLAG beads (Merck Millipore) in buffer 1. Following an 18h-incubation at 4°C, the protein was eluted by adding 500 μg/ml of 3xFLAG peptide (China Peptides CO LTD) in buffer 2 (50 mM HEPES pH 7.5, 500 mM KCl, 10 mM MgCl_2_, 10% glycerol and 0.5 mM TCEP).

Bacteria expressing PolγB were lysed by sonication (10 min; 3 s ON/6 s OFF cycles at 37% amplitude) in buffer 3 (50 mM HEPES pH 7.5, 500 mM KCl, 0.1 mM EDTA, 10 mM Imidazole, 10% glycerol, 2 mM β-mercaptoethanol, and cOmplete™ Protease Inhibitor Cocktail). The clarified supernatant (50,000 x *g* for 45 min) was loaded onto a HisTrap column (Cytiva), and the protein was eluted in buffer 3 supplemented with 500 mM imidazole. The eluate was dialysed overnight against buffer 4 (50 mM HEPES pH 7.5, 400 mM KCl, 10% glycerol, 10 mM MgSO_4_ and 0.5 mM TCEP) in the presence of SUMO protease to remove purification tags. The tag-free PolγB was collected in the flow-through of a second HisTrap purification step. A final polishing step was performed using size-exclusion chromatography (Superdex 200 10/300 GL, Cytiva) in buffer 4. Fractions containing PolγB were concentrated using a 30 kDa MWCO Amicon Ultra-15 centrifugal filter device (Merck) and stored at −80°C.

Bacteria expressing mtSSB were lysed by sonication (10 min; 3 s ON/6 s OFF cycles at 37% amplitude) in buffer 5 (20 mM Tris pH 6.8, 1 M KCl, 10 mM MgCl_2_, 5% glycerol, and cOmplete™ Protease Inhibitor Cocktail). After centrifugation at 50,000 x *g* for 45 min, the soluble fraction was loaded onto a HisTrap column, and the protein was eluted using buffer 6 (20 mM HEPES pH 8, 150 mM KCl, 10 mM MgCl_2_, 5% glycerol and 500 mM imidazole). The eluate was then dialysed overnight at 15°C against buffer 6 without imidazole in the presence of PreScission protease to remove the Strep tag. The protein was concentrated, and glycerol was added to a final concentration of 15% before storage at −80 °C.

Bacteria expressing PolγA-catcher fused to MBP were lysed by sonication (5 min; 3 s ON/1 s OFF cycles at 37% amplitude) in buffer 7 (50 mM HEPES pH 7.5, 500 mM KCl, 5% glycerol, 1 mM β-mercaptoethanol and cOmplete™ Protease Inhibitor Cocktail). After centrifugation at 50,000 x *g* for 45 min, the soluble fraction was loaded onto a HisTrap column, and the protein was eluted with buffer 8 (50 mM HEPES pH7.5, 300 mM KCl, 5% glycerol, 500mM Imidazole and 1 mM β-mercaptoethanol). The protein was concentrated, and a final polishing step was performed using size exclusion chromatography (Superdex^TM^ 75 10/300 GL, Cytiva) in buffer 9 (50 mM Tris pH 7.5, 150 mM KCl, 5% glycerol, 1 mM β-mercaptoethanol). The protein was concentrated before storage at −80 °C.

### 4.4. Polγ holoenzyme reconstitution

To reconstitute Polγ holoenzyme, wild-type or mutant versions of PolγA and PolγB were mixed at a final concentration of 5-10 μM in buffer 10 (50 mM Tris pH 7.5, 250 mM KCl, 10 mM MgCl2 and 1 mM TCEP) with a 1.2-fold molar excess of PolγB (calculated as a dimer), and incubated on ice for 10 min. Next, analytical gel filtration chromatography was performed using a Superdex 200 Increase 3.2/300 column (Cytiva) on an ÄKTA pure micro system. The fractions containing Polγ holoenzyme were pooled, concentrated, flash-frozen and stored at −80 °C for later use in biochemical assays or directly for EM grid preparation.

### 4.5. DNA substrates

DNA and RNA substrates used in primer extension and strand-displacement DNA replication assays, fluorescence anisotropy experiments, and cryo-EM sample preparation were purchased from IDT and Merck. When annealing was required, samples were incubated at 95 °C for 5 min, followed by gradual cooling at a rate of 1 °C/min using a thermocycler. A detailed list of primers and DNA substrates is provided in Supplementary Table 2. G4 DNA substrates were prepared as described in *et al*. 2020^21^.

DNA substrates for single-molecule studies were prepared as described before^47,61,62^. Briefly, for primer extension assays, we used a gapped DNA template consisting of ~900 nt of ssDNA flanked by ~3550 bp dsDNA handles. The DNA construct was prepared from the *pBacgus*11 vector (Novagen) and labelled with digoxigenin (Dig) at one end and biotin at the other. The approximate length of the ss-DNA fragment was determined by denaturing gel electrophoresis. For strand displacement activities, the hairpin DNA construct consisted of a 2686 base pairs (bp) DNA ‘handle’ (pUC19 vector, Novagen) labelled with Dig at one end, a 5’ (dT)_35_ end functionalised with biotin, and a 559 bp stem capped by a (dT)_4_ loop. The final hairpin construct contains a unique 3’ end loading site for the DNA polymerase.

### 4.6. Negative stain (NS-EM) sample preparation and data acquisition

NS-EM samples were prepared by incubating 5 μL of freshly reconstituted protein complexes at 0.05 mg/mL on glow-discharged (Quorum GloQube glow discharger: 15 mA, 45 s, 0.1 bar) carbon grids (Electron Microscopy Sciences, CF400-Cu) for 45 s. After removing the excess of sample, grids were washed three times with 5 μL of ultrapure H_2_O and fixed with 5 μL of 1 % uranyl formate for 30 s. Grids were air-dried and stored at RT for later use. NS-EM data were acquired on an in-house FEI Tecnai G2 Spirit microscope operated at 120 kV with a TVIPS TemCam-F416R camera. Images were collected with a total dose of 30 e^−^/Å^2^ and a defocus range of 0.6-1.4 μm at 42.000x nominal magnification (calibrated pixel size of 2.5 Å).

### 4.7. Chemical grid pre-conditioning for Cryo-EM

Quantifoil R2/1 mesh 300 grids were incubated for 10 s in a DMSO solution containing 5 nM 1-Pyrenebutyric acid (Merck; ref. 257354). The grids were then washed sequentially in 100% isopropanol and in 100% ethanol for 10 s each. After washing, the grids were air-dried for at least 2 h and stored until firther use. Grids prepared using this method did not require plasma treatment and could be stored at RT for up to one week before use.

### 4.8. Cryo-EM sample preparation and data acquisition

For cryo-EM sample preparation, freshly reconstituted Polγ complexes were used, as described above. In the Polγ-mtSSB-DNA dataset, Polγ complex fractions were pooled, diluted to 0.8 μM and mixed with 1 μM DNA (60mer-ssDNA), 1 μM mtSSB (tetramer) and 1 mM ddATP. After a 10 min incubation on ice, 3 μL of the reconstituted complex was deposited onto Quantifoil grids (R 0.6/1, 300 mesh) that had been positively charged using a Quorum GloQube glow discharge device and Amylamine vapour (TCI EUROPE, TCIAA0446) for 45 s at 15 mA and 0.1 bar pressure. The excess sample was then blotted for 3.5 s, and the grids were plunge-frozen using a VitroBot Mark III (FEI) with force −14 at 4 °C and 99 % humidity. In the Polγ-sdDNA dataset, Polγ complex fractions were pooled, diluted to 1 μM and mixed with 2 μM DNA (sdDNA). DNA polymerisation was initiated by adding dNTPs to a final concentration of 1 mM. Immediately after initiating the reaction, 3.5 μL of the mixture was deposited onto chemically pre-conditioned grids. The excess sample was blotted for 2 s, and the grids were plunge-frozen using a VitroBot Mark III (FEI) with force −16 at 4 °C and 99 % humidity. In the Polγ-G4 dataset, Polγ complex (exonuclease-deficient PolγA) fractions were pooled, diluted to 0.8 μM and mixed with 1.6 μM DNA (G4: 656-1056) in the presence of dCTP, dGTP, dTTP and ddATP (to block the unwrapping of the G4). The reaction was incubated at 20°C for 5 minutes. Later, 3.5 μL of the mixture was deposited onto chemically pre-conditioned grids.

Initial screening of sample preparation conditions was performed on an in-house FEI Tecnai G2 Spirit microscope with a TVIPS TemCam-F416R camera and on a JEM-2200FS (JEOL) equipped with a Gatan K3 direct electron detector. After best conditions were identified, complete datasets were collected on a FEI Titan Krios operated at 300 kV and equipped with a Gatan K3 direct electron detector, either at eBIC (Electron Bio-Imaging Centre, Diamond Light Source, UK) or BREM (Basque Resource for Electron Microscopy, Biofisika Institute, Spain). For details on data collection conditions see Table 1.

### 4.9. EM data processing

Data processing followed a similar scheme across all datasets, using RELION-5^63^. Briefly, data processing started with motion correction using the MotionCor2 CPU version^64^ and CTF parameters estimated using CTFFind-4.1^65^, both implemented in RELION-5. Micrographs were examined after CTF and ice thickness estimation, and those with extreme attributes were removed using relion_analyse.py for all datasets^66^.

A total of 29.117 micrographs from the initial 30,577 were used to determine the structure of Polγ-DNA-mtSSB. The dataset showed high homogeneity and after picking and cleaning up the dataset using 2D and 3D classification in Relion4 and CryoDRGN^67^, a selection of 638,328 particles was used for the final reconstruction. After initial refinement using gold-standard protocols in RELION-5, we obtained a map at 3.08Å resolution (FSC 0.143 cutoff criteria). CTF-refinement and Bayesian polishing applying pre-multiplied CTF in Relion improved the map and the resolution of the reconstruction to 2.47 Å. We ran focused refinements on PolγA, PolγB using different masks centered on the different features. With this approach, we obtained more defined maps of these regions, reaching resolutions of 2.35 Å and 2.93 Å. For the resolution of the catcher domain, the particles were re-extracted so the center of mass of the re-extracted particles was the catcher. After 3D curation and refinement, the cather-centered volume was resolved at 3.17 Å, from which we could model it (see Supplementary Fig. 3 for details).

A total of 23,097 micrographs from the initial 29,054 were used to determine the structure of Polγ-SD-DNA. After picking and cleaning up the dataset using 2D and 3D classification in RELION-4.0^68^, three populations of particles were identified: Polγ-APO (984,666 particles), Polγ-SD-DNA-open (225,443 particles), and Polγ-SD-DNA-closed (836,840 particles). Each individual subset of particles was refined, yielding reconstructions at 3.25 Å, 3.37 Å and 3.00 Å, respectively. For the Polγ-DNA-mtSSB dataset, we performed focused refinements on PolγA and PolγB, improving resolution in all cases. For Polγ-APO, focused refinement on PolγA and PolγB yielded resolutions of 3.00 Å and 3.29 Å, respectively. For Polγ-SD-DNA-open, resolutions of 3.25 Å and 3.21 Å were obtained for PolγA and PolγB, respectively, while for Polγ-SD-DNA-closed, the corresponding resolutions were 2.88 Å and 2.90 Å. (see Supplementary Fig. 1 for details).

A total of 48,812 micrographs were used to determine the structure of Polγ-G4-DNA, comprising 24,109 acquired on the flat stage and 24,703 on the 30° tilted stage. After picking and cleaning the dataset using 2D and 3D classification in RELION-5 and cryoDRGN, four particle populations were identified: Polγ-G4-1 (209,158 particles), Polγ-G4-2 (185,759 particles), Polγ-G4-3 (132,600 particles), and Polγ-G4-4 (68,861 particles). Each individual subset of particles was subsequently refined, yielding reconstructions at 3.22 Å, 3.19 Å, 3.33 Å and 3.41 Å, respectively. As for the previous datasets, focused refinements on PolγA, PolγB were performed, improving the resolution in all the cases. For Polγ-G4-1, focused refinement on PolγA and PolγB yielded resolutions of 3.05 Å and 3.37 Å, respectively. For Polγ-G4-2, resolutions of 3.05 Å and 3.29 Å were obtained for PolγA and PolγB, respectively. For Polγ-G4-3, the corresponding resolutions were 3.19 Å and 3.41 Å, while for Polγ-G4-4, they were 3.25 Å and 3.37 Å, respectively (see Supplementary Fig. 2 for details).

### 4.10. Model building and analysis

We fitted the preexisting X-ray model 5C52^37^ as a rigid body into the cryo-EM maps using MOLREP^69^. We also did a *de novo* modelling using Modelangelo^70^ implemented in RELION-4.1 and AlphaFold3^71^ predictions to model the regions that were unresolved in the previous structures. Further model building was performed in Coot and ISOLDE using sharpened maps and assisted by denoised maps obtained with EMReady2. Composite maps were generated in UCSF ChimeraX^72^ by combining the consensus reconstruction with locally refined maps obtained from focused refinements (e.g., PolγA, PolγB, and catcher-centred refinements), improving visualisation of regions exhibiting different local resolutions. These processed and composite maps were used exclusively to assist model building, interpretation, and visualisation, whereas all refinements were carried out against the corresponding experimental half maps using REFMAC5-Servalcat^73^ within the CCPEM^74^ suite with automatic weighting, and PHENIX real space refinement^75^.

The visual analysis and generation of figures were performed using UCSF ChimeraX^72^ and PyMol^76^. Protein interfaces and intermolecular interactions were analysed using the PDBePISA^77^ server.

In the Polγ-APO structure, the high flexibility of the PolγB dimer prevented accurate modelling of the protein. Therefore, the PolγB dimer model from the Polγ-DNA-mtSSB structure was fitted into the corresponding density in the Polγ-APO map. Similarly, the density corresponding to the distal PolγB subunit in the Polγ-SD structures did not allow reliable model building. Consequently, only the distal PolγB subunit from the Polγ-DNA-mtSSB structure was fitted into these densities, meanwhile the proximal PolγB was properly modelled.

### 4.11. Primer extension and strand displacement assays

DNA synthesis was evaluated *in vitro* using synthetic substrates (see supplementary Table 2) designed to report PE in the absence of strand displacement when only dTTP and dCTP are provided, and SD activity upon addition of all four dNTPs (Fig. 3b). The reconstituted Polγ complex (20 nM) was incubated with a substrate (20 nM) containing either an IRDye700-labelled primer for gel-based analysis or a FAM-labelled primer for capillary electrophoresis analysis. Reactions were conducted at 37 °C in a buffer containing 50 mM Tris pH 7.5, 50 mM KCl, 10 mM MgSO_4_, 0.1 mg/ml BSA, 10% glycerol, 2 mM DTT and 200 μM dNTPs for strand displacement DNA synthesis or 200 μM dTTP and dCTP for the primer extension polymerase assay. Reactions were stopped at 10 and 30 min by adding an equal volume of a stop buffer containing 90% formamide, 50 mM EDTA, and 0.04% (w/v) bromophenol blue, followed by heating at 95 °C for 5 min. Polγ-Δ991-1051 was assayed at a 6fold higher concentration because of its lower basal polymerase activity.

Reaction products were analysed either by capillary electrophoresis (see below) or by gel electrophoresis in 15% acrylamide TBE-urea gels (Invitrogen). Gels were run at 180 V for 50 min in 1x TBE buffer. Fluorescently labelled products were visualised using an Odyssey CLx Imager (LI-COR Biosciences). For each independent experiment, the ratio between strand-displacement and primer-extension activity was calculated. Each Polγ variant was compared with wild-type Polγ using a two-tailed paired Student’s t-test, with measurements paired by experimental replicate.

G4 traversal was evaluated using primer-template substrates containing the G4a or G4b sequence (Supplementary Table 2). Reactions contained all four dNTPs and were performed under the same conditions as described above. Polγ-Δ991-1051 was assayed at a 6fold higher concentration because of its lower basal polymerase activity. G4 traversal was quantified from fluorescence intensity profiles of the electrophoresis lanes. For each reaction, signal was integrated separately within two predefined regions corresponding to products terminating before or within the G-rich sequence and products extending beyond it, as indicated in Fig. 5a. The unextended primer was excluded from the stalled-product signal. The processed-to-stalled product ratio was calculated by dividing the integrated intensity of products extending beyond the G-rich sequence by that of products terminating before or within this region.

### 4.12. Capillary Electrophoresis

Reaction products were also analysed using capillary electrophoresis in a 3730xl Genetic Analyzer (Applied Biosystems). Data were processed using custom software (https://github.com/cryoEM-CNIO/CE_tools) to extract and quantify fluorescence intensity values, after migration time and intensity normalisation with an internal loading and migration control included in all reactions (4 nM FAM-labelled DNA added to the stop buffer; see control-DNA in Supplementary Table 2). For each independent experiment, the ratio of SD to PE activity was calculated. Each Polγ variant was compared with wild-type Polγ using a two-tailed paired Student’s t-test, with measurements paired by experimental replicate.

### 4.13. Fluorescence Polarisation assay

Fluorescence Polarisation (FP) assays were carried out in 384-well low-volume round-bottom plates (Corning) at 25 °C using 6-FAM labelled primer-template DNA substrates (see Supplementary Table 2) on a CLARIOstar Plus plate reader (BMG LABTECH GmbH, Germany). Fluorescence was excited at a wavelength of 482 nm (±16), and polarisation was detected at 530 nm (±40). FP values were calculated using the CLARIOstar software and analysed using GraphPad Prism 11 (GraphPad software Inc., USA). Each experiment was performed in at least three independent replicates.

For MBP-catcher samples, serial protein dilutions were prepared in a buffer containing 50 mM HEPES pH 7.5, 25 mM KCl, and 5% glycerol, with final protein concentrations ranging from 448 μM to 13 nM. Samples were mixed with DNA substrates at a constant concentration of 10 nM. Full-length PolγA measurements followed a similar protocol, using a buffer containing 50 mM HEPES pH7.5, 50 mM KCl, 10 mM MgCl2, 1 mM DTT and 5% glycerol and with initial maximum concentrations ranging from 4 μM to 9.6 μM.

For each independent experiment, FP values were normalised to the maximum response within that experiment and fitted separately in GraphPad Prism using the “agonist versus normalised response - variable slope” model to account for the sigmoidal binding behaviour observed experimentally. We obtained apparent KD values from each independent fit and report them as mean ± SD from at least three independent experiments; exact n values are provided in the corresponding figure legends. For graphical representation, we combined normalised data from the independent experiments and fitted them globally using the same model.

### 4.14. Protein thermal stability

The thermal stability of PolγA (WT and variants) was assessed using a Tycho NT.6 instrument (NanoTemper Technologies). Samples were prepared at a concentration of 2.5 μM in buffer containing 50 mM HEPES pH 7.5, 500 mM KCl, 10 mM MgCl_2_, 10% glycerol and 0.5 mM TCEP. For each measurement, 10 μL of the protein solution was loaded into capillaries supplied by the manufacturer. The thermal unfolding profile was recorded by heating the sample from 35 °C to 95 °C at a rate of 30 °C/min. Changes in the fluorescence ratio (350 nm/330 nm) were monitored to detect shifts in protein structure during heating. The inflection point of the fluorescence ratio (Tm) was calculated using the Tycho NT.6 software. All measurements were performed in duplicate or triplicate.

### 4.15. Molecular Dynamics (MD) simulations

Three MD simulations were conducted for each Polγ mutant as well as for the wild type protein using GROMACS 2024.1^78^ with AMBER99SB force field^79^ and TIP3P water models^80^, fully solvating the simulation boxes with water and 150mM NaCl, neutralising the system. The simulations were restricted to the residues of the displacement loop motif (from residue 991 to residue 1052). The studied mutants included R1006A, E1016A, R1026A, R1026A-R1030A-R1034A (RRR), R1030A, R1034A, W1038A, and W1041A. Long-range electrostatic interactions were calculated using the Particle Mesh Ewald (PME)^81^ algorithm with a cutoff of 1.2 nm. All structures were subject to energy minimisation using the steepest descent method until reaching a Fmax < 10 kJ mol^−1^ nm^−1^, with a step size of 0.1 Å and updating the neighbour list at each step. A 500 ps NVT equilibration was performed using a velocity rescale thermostat^82^, followed by a 1 ns NPT equilibration using a Parrinello-Rahman^83^ barostat, both with a timestep of 2 fs. The production runs lasted 50 ns with a timestep of 2 fs. Stability of the proteins was assessed by calculating the Root Mean Square Deviation (RMSD) and the radius of gyration of the protein backbone, using the gmx_rms and gmx_gyrate utilities included in GROMACS 2024.1.

### 4.16. Single-molecule force spectroscopy

A counter-propagating dual-beam optical tweezers instrument^84^ was used to manipulate individual DNA constructs, a primer-template DNA molecule (for PE) or fork DNA structure (for SD), functionalised with biotin or digoxigenin at each end. The DNA was tethered between streptavidin- and anti-digoxigenin-coated polystyrene beads (3 μm) between the beads on top of a micropipette and in the optical trap, respectively. Polγ and SSB proteins were diluted to concentrations of 2 and 5 nM, respectively, in the reaction buffer and subsequently injected into the flow cell. Data was monitored at 5100 Hz at 22±1 °C using a feedback loop to maintain a constant mechanical tension on the DNA construct. The trap stiffness was k=0.135±0.0043 pN nm^−1^ (3.0 μm beads). With this setup, as the polymerase unwinds the DNA fork (SD) or converts ssDNA to dsDNA (PE), it changes the distance between the beads or the tether length (Fig. 4).

### 4.17. Single-molecule data analysis

The number of nucleotides replicated in the strand-displacement assays was calculated from the increase in tether extension. This extension change was divided by the increase in molecular length, at the applied force, associated with each catalytic step. Each step generates one newly synthesised base pair and one displaced single-stranded nucleotide. The number of nucleotides incorporated in primer extension assays was obtained by dividing the change in tether extension by the change in extension due to the conversion of one single-stranded nucleotide into its double-stranded counterpart at a given force^85,86^. The extension of the dsDNA was approximated with the worm-like chain model for polymer elasticity with a persistent length of P = 53 nm and stretch modulus S = 1200 pN/ nm^87^. The average extensions of SSB-free and SSB-bound ssDNA nucleotides as a function of mechanical tension were reported previously^18^.

The average replication rates at each force *V* _mean_(*f*) were determined by a line fit to the traces showing the number of replicated nucleotides versus time. The final rate at each tension was obtained by averaging over all of the traces taken at similar tension values (±1 pN). The average replication rate without pauses at each tension, *V* (*f*), was determined with an algorithm that computes the instantaneous velocities of the trajectory, averaging the position of the holoenzyme along the DNA over sliding time windows, as described previously^18^. The average residence time at the pause state per nucleotide at each tension, *T*_*p*_(*f*), was calculated as the difference between the average total residence time per nucleotide (*T*_*t*_(*f*)= 1/ *V* _mean_(*f*)) and the residence time in the active state (*T*_*a*_(*f*)= 1/ *V* (*f*)).

## 5. ACKNOWLEDGEMENTS

This study was supported by MCIN/AEI/10.13039/501100011033 (grants BFU2017-87316-P, CNS2023-143762, and PID2024-159675NB-I00 to RFL; PID2021-126755NB-I00 and PID2024-158478NB-I00 to BI; PRE2019-088885 to I.P.G.A.), by La Caixa Foundation (HR24-00604) to RFL and BI, and the National Institute of General Medical Sciences of the National Institutes of Health under Award GM139104 to G.L.C. We acknowledge the help of Jasminka Boskovic and Johanne Lecoq from the CNIO EM facility for all their help on the cryoEM experiments, as well as Luís Lombardía Ferreira and Diana Romero Gómez from the CNIO Molecular Diagnostics Unit for all their help with the capillary electrophoresis experiments. Cryo-EM data was obtained at the Diamond Light Source cryo-EM facility at the UK’s National Electron Bio-imaging Center (eBIC) under BAG “Stop cancer-structural studies of macromolecular complexes involved in cancer by cryo-EM” (BI39322-1) and at the Basque Resource for Electron Microscopy located at Instituto Biofisika (UPV/EHU, CSIC), supported by the Department of Science, Universities and Innovation and the Innovation Fund of the Basque Government, with additional support from MCIN (Recovery, Transformation and Resilience Plan) and the Basque Government “Biotechnology Complementary Plan Applied to Health” with funding from European Union NextGenerationEU [PRTR-C17.I1, PRTR-C17.I01.P01.S13, AAAA_ACG_AY_2539/22_05].

## 6. AUTHOR CONTRIBUTION

S.M.-A. and A.G.-G. contributed equally to this work. S.M.-A. purified proteins, reconstituted protein complexes, performed cryo-EM data acquisition, processing and analysis, carried out biochemical and biophysical experiments, contributed to the study conceptualisation, and prepared the first draft of the manuscript. A.G.-G. purified proteins, reconstituted protein complexes, and performed the biochemical experiments. I.P.G.A. performed and analysed the single-molecule experiments. E.C.-P. purified proteins and reconstituted protein complexes. E.A.-U. performed the evolutionary analyses. M.M.-M. contributed to cryo-EM data analysis and revised the manuscript. C.C.-S. performed the molecular dynamics simulations. G.L.C. contributed to data interpretation and revised the manuscript. B.I. supervised the single-molecule studies, secured funding, and revised the manuscript. R.F.-L. conceived and supervised the study, secured funding, performed cryo-EM data acquisition, processing and analysis, and wrote and edited the manuscript. All authors discussed the results and reviewed and approved the final manuscript.

## 7. COMPETING INTERESTS

The authors declare no competing financial interests.

## 8. DATA AND MATERIALS AVAILABILITY

Materials are available upon request to R. F.-L., subject to standard material transfer agreements. Cryo-EM maps and atomic models have been deposited in the Electron Microscopy Data Bank (EMDB) and the Protein Data Bank (PDB) under accession codes EMD-XXXXX and XXXX. The entries are currently on hold and will be released upon publication; until then, the maps and models can be requested by email from the corresponding author.

## 9. CODE AVAILABILITY

The custom software used to process and quantify capillary electrophoresis data, including migration-time and fluorescence-intensity normalisation, is publicly available at https://github.com/cryoEM-CNIO/CE_tools.

## Notes

### Competing Interest Statement

The authors have declared no competing interest.

### Summary of Updates

Updated data availability and table S1

