## Supplementary Figures and Tables for "A dedicated motif in human polymerase gamma enables DNA synthesis through replication roadblocks"


Supplementary figures and tables for the accompanying manuscript. Typeset with the rho-class LaTeX template.

### 1. SUPPLEMENTARY FIGURES

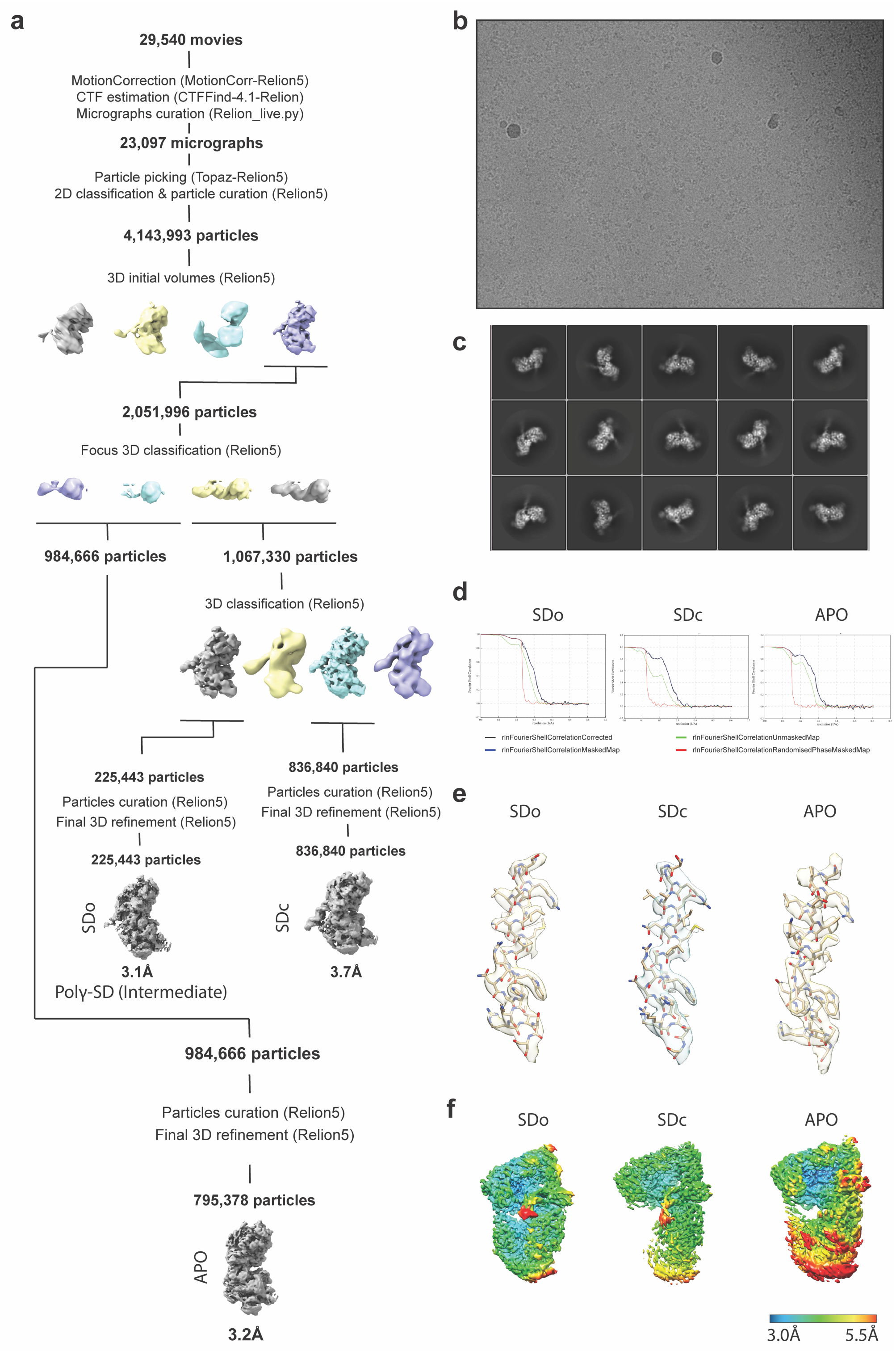


**Supplementary Figure 1. Cryo-EM processing of the Polγ-SD dataset. a.** cryo-EM processing workflow for Polγ bound to the strand-displacement DNA substrate. Particle classification identified a DNA-free Polγ population and two DNA-bound conformations corresponding to the open and closed strand-displacement states. Subsequent 3D refinement and focused refinements on PolγA and PolγB improved the corresponding reconstructions. Representative densities illustrating the alternative DNA-bound conformations are shown together with local-resolution maps. **b.** Representative micrograph. **c.** Representative 2D class averages. **d.** Fourier shell correlation curves for the final reconstructions. **e.** Details of the thumb helix density in each state. **f.** Local resolutions of the final reconstructions.

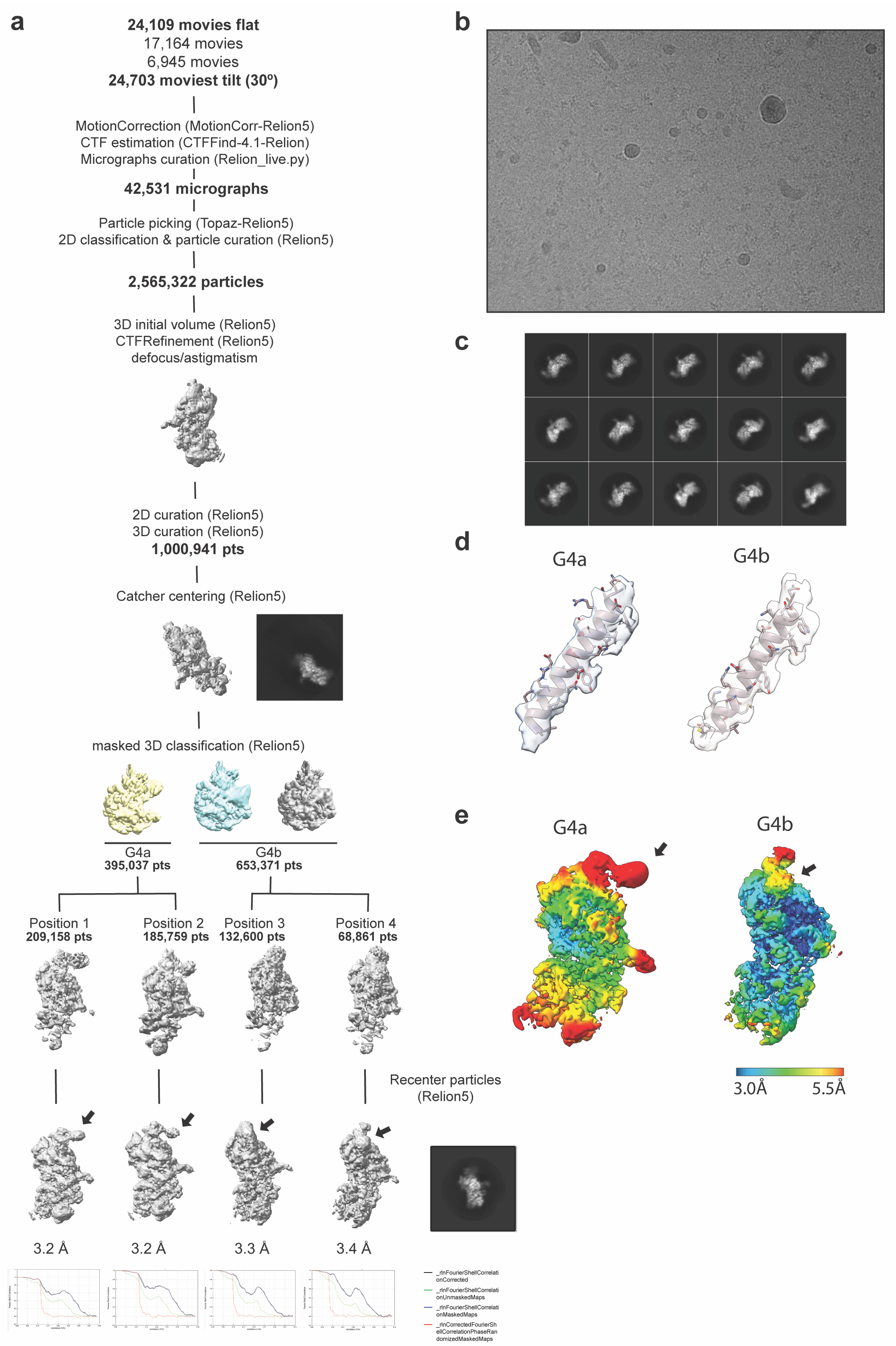


**Supplementary Figure 2. Cryo-EM processing of the Pol$\gamma$-G4 DNA dataset.** a. processing workflow for Pol$\gamma$ bound to G4-containing template DNA. Data acquired at 0° and 30° stage tilt were combined for processing. Following particle picking, 2D classification and initial 3D refinement, particles were subjected to further classification to resolve alternative positions of the G4-containing DNA substrate. Four major structural classes were obtained and independently recentred and refined. Fourier shell correlation curves for the final reconstructions are displayed. **b.** Representative micrograph. **c**. Representative 2D class averages. **d.** Details of the thumb helix density in each state. **e.** Local resolutions of the final reconstructions.

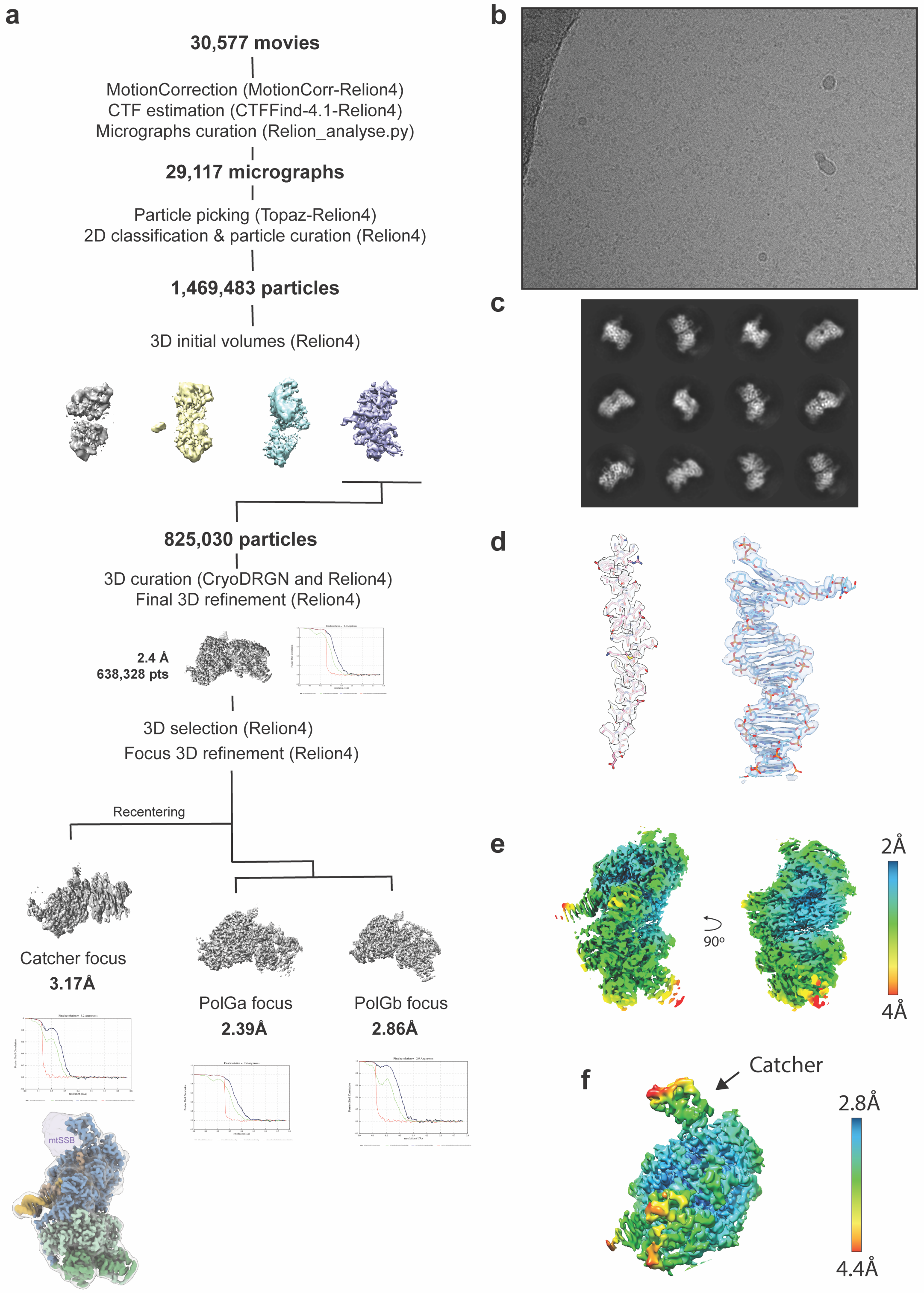


**Supplementary Figure 3. Cryo-EM processing of the Polγ-DNA-mtSSB dataset. a.** Scheme of the cryo-EM processing strategy used to obtain the Polγ–DNA–mtSSB reconstruction. Following motion correction, CTF estimation, particle picking and 2D classification, particles were subjected to successive 3D classification and refinement steps. Focused refinements centred on PolγA and PolγB improved the corresponding local densities, while recentering particles on the DNA-engaging region enabled improved visualisation of the flexible template-engaging element. Fourier shell correlation curves for the final reconstructions are displayed **b.** Representative micrograph. **c.** Representative 2D class averages. **d.** Details of the thumb helix and DNA densities. **e.** Local resolutions of the final reconstructions. **f.** Local resolution of PolγA centered reconstruction showing the catcher-domain density.

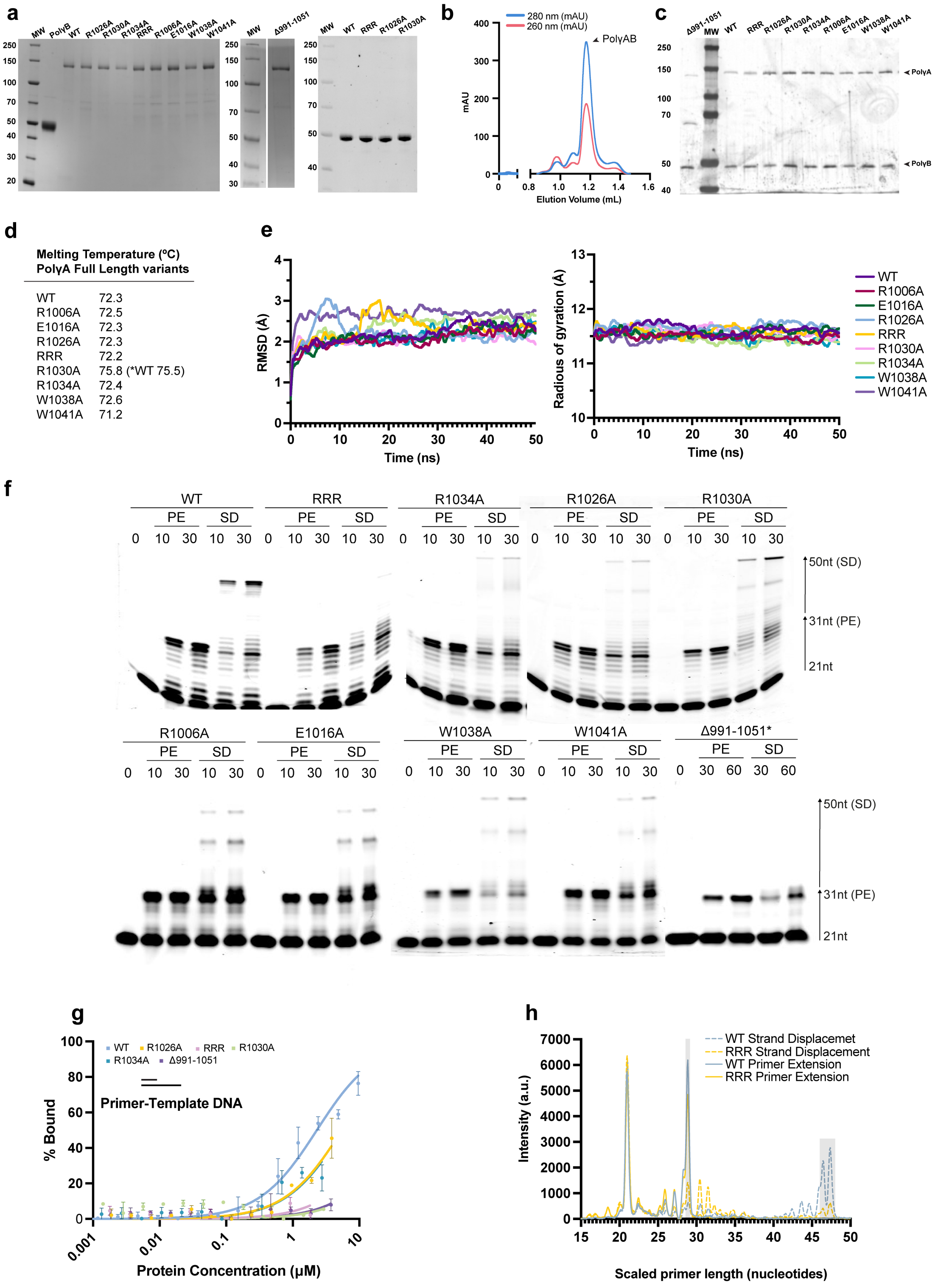


**Supplementary Figure 4. Purification, stability and functional characterisation of Polγ variants. a,** SDS–PAGE analysis of purified PolγA full-length or truncated (MBP-991-1051) variants (left/middle gels and right gel, respectively) used in this study. **b,** Representative size-exclusion chromatography profile of reconstituted Polγ holoenzyme. **c,** SDS-PAGE analysis of reconstituted Polγ variants. **d,** Apparent thermal-transition temperatures measured for WT PolγA and the indicated variants. **e,** Molecular-dynamics analysis of the isolated 991-1051 constructs. Root-mean-square deviation (RMSD; left) and radius of gyration (right) are shown for WT and mutant proteins during the simulations. **f,** Primer-extension (PE) and strand-displacement (SD) assays for the extended panel of Polγ variants at the indicated reaction times. Asterisk indicates protein concentrations 6-fold higher than those used for the other proteins. **g,** Fluorescence-polarisation analysis of DNA binding by full-length WT and mutant Polγ proteins using primer-template DNA. **h,** Representative capillary-electrophoresis traces showing products generated during primer-extension and strand-displacement reactions by WT and RRR Polγ. Grey boxes indicate regions used for quantification.

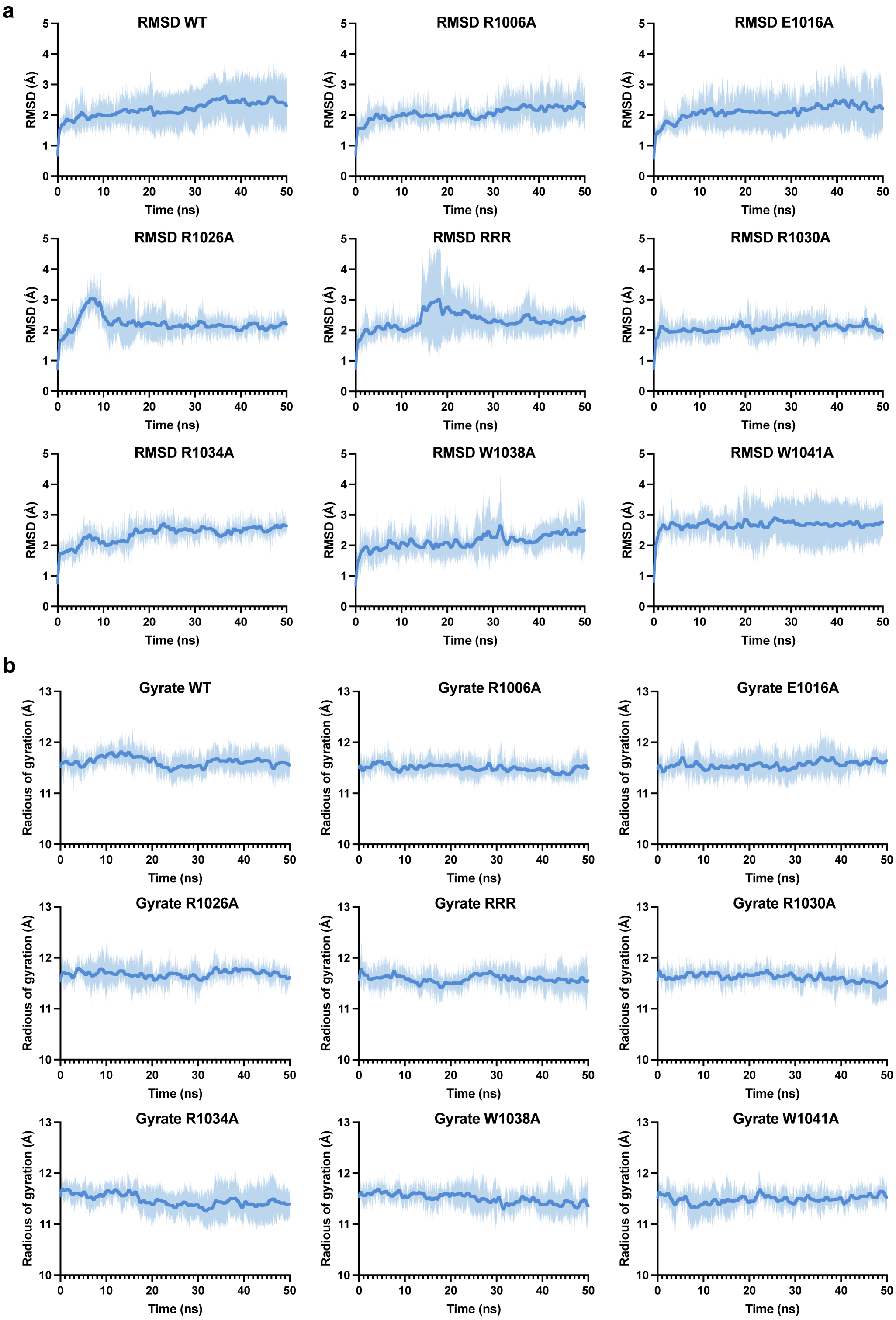


**Supplementary Figure 5**. **Molecular-dynamics analysis of WT and mutant Pol$\gamma$ template-engaging elements. a,** Root-mean-square deviation (RMSD) of the catcher backbone during 50-ns molecular-dynamics simulations of WT and the indicated Pol$\gamma$ variants. **b,** Radius of gyration calculated over the same simulations. Lines represent the average over 3 simulations; shaded regions represent the standard deviation of the simulations.

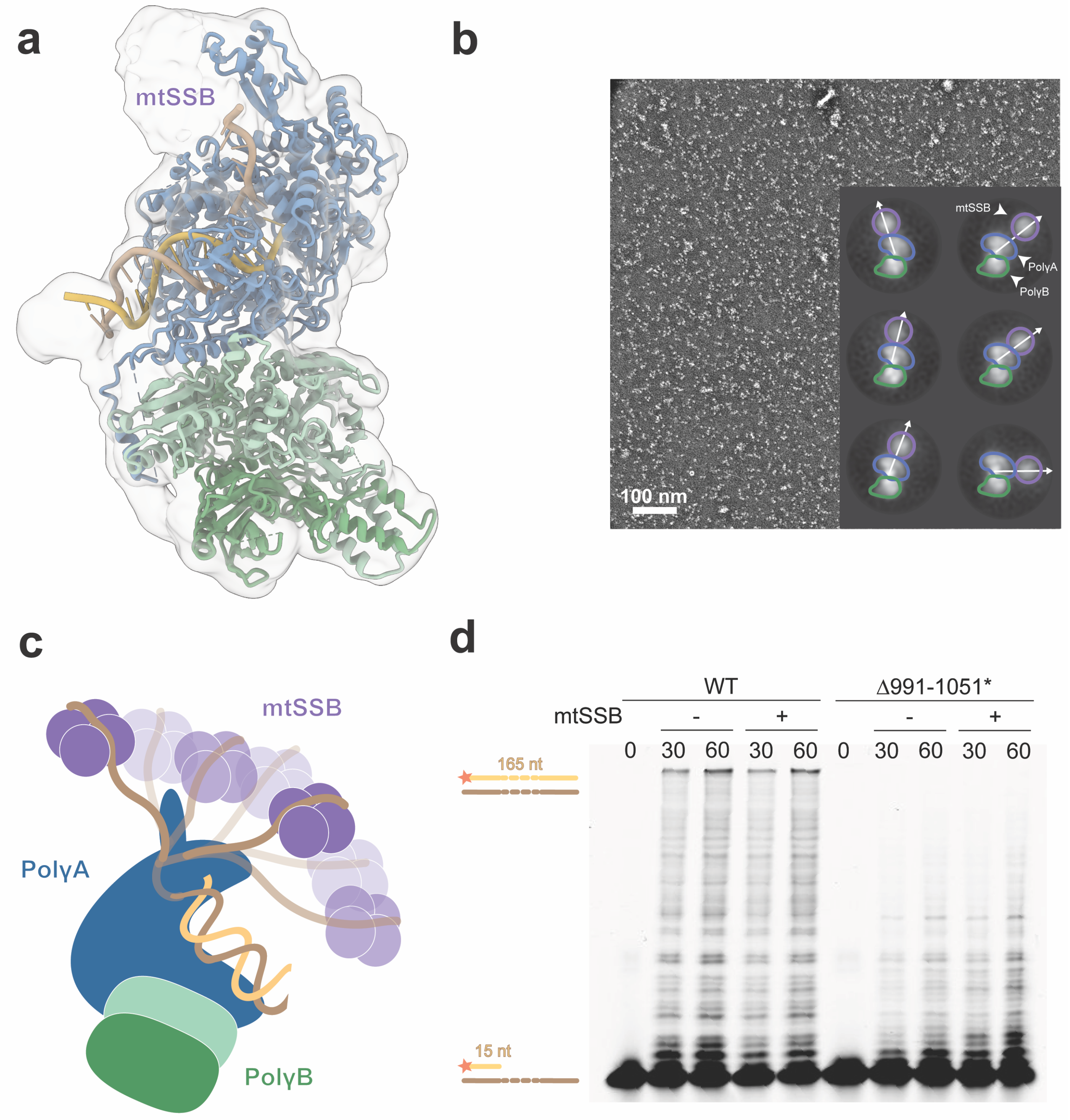


**Supplementary Figure 6. Structural and functional analysis of Polγ in the presence of mtSSB. a,** Cryo-EM reconstruction of Polγ bound to DNA in the presence of mtSSB, shown at a threshold that reveals additional low-resolution density compatible with mtSSB adjacent to the incoming ssDNA. **b,** Representative negative-stain electron micrograph of the Polγ-DNA-mtSSB complex and selected 2D class averages. Alignment on Polγ reveals heterogeneous positions of the additional mtSSB density relative to the polymerase. **c,** Schematic representation of the range of positions sampled by mtSSB relative to Polγ and the incoming ssDNA. **d,** Primer-extension assays performed with WT Polγ and Polγ-Δ991–1051 on a long ssDNA template in the absence or presence of mtSSB for the indicated reaction times. Asterisk indicates protein concentrations 6-fold higher than those used for the other proteins.

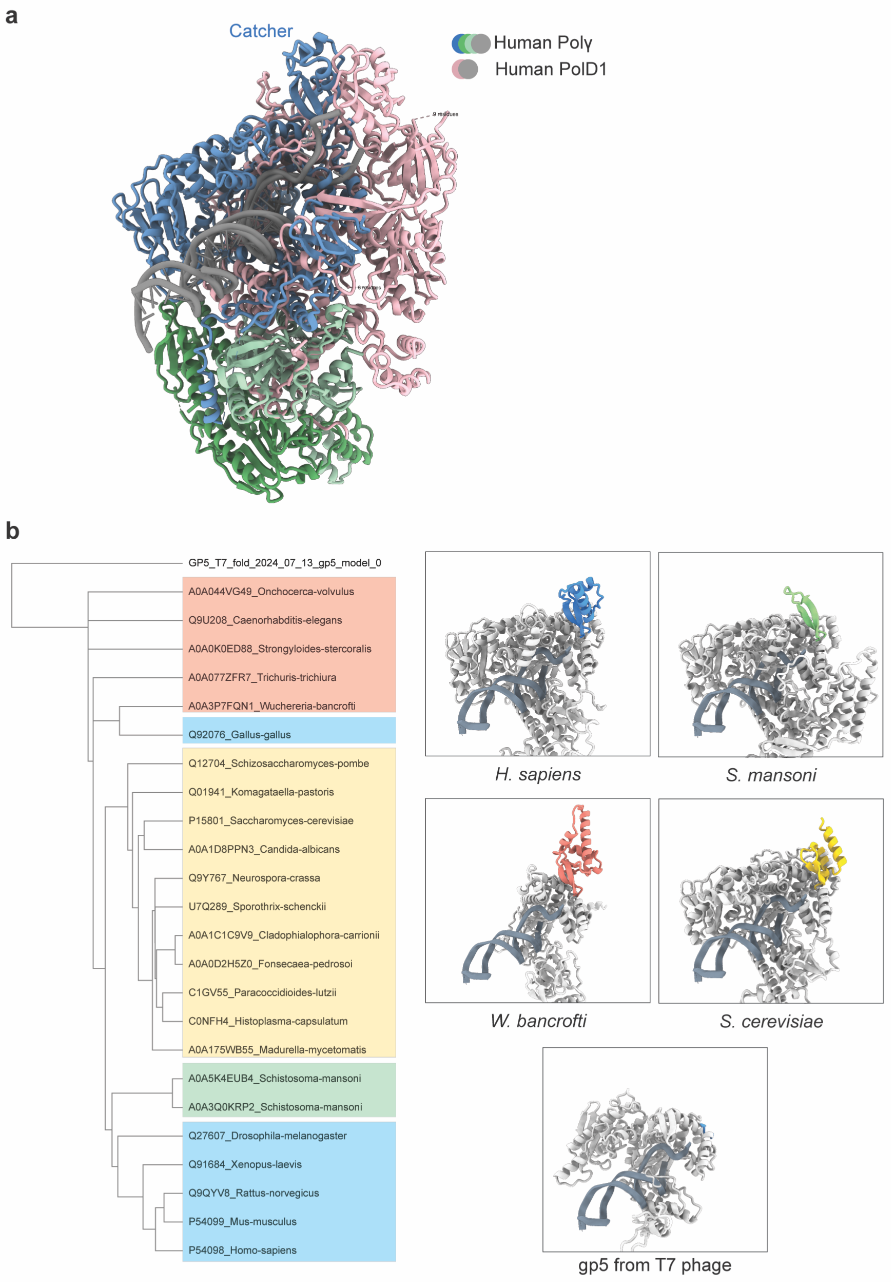


**Supplementary Figure 7. Structural and evolutionary comparison of catcher-like elements. a**, structural superposition of human Polγ and human Polδ catalytic subunit (*POLD1*), highlighting the catcher of Polγ and the corresponding insertion in Polδ positioned near the DNA entry channel. Structures were aligned based on DNA position. **b,** Left, phylogenetic representation of the structural analysis performed with FoldSeek and FoldMason. Major groups are distinguished by background shading. Right, structural comparison of DNA polymerases from different species, highlighting insertions positioned adjacent to the DNA entry channel. The conserved polymerase core is shown in grey, while the corresponding insertion in each structure is coloured according to its phylogenetic group.

**a**

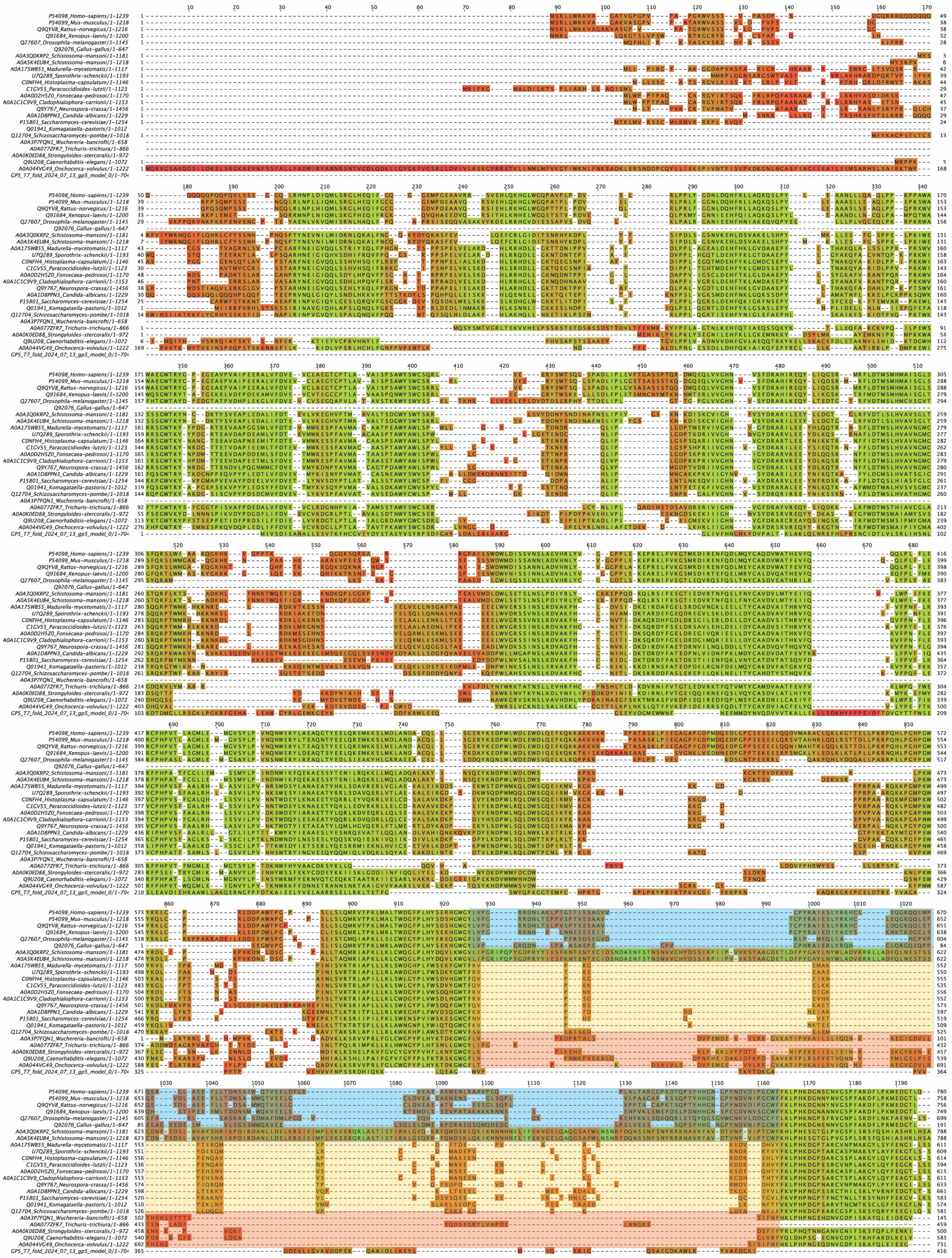

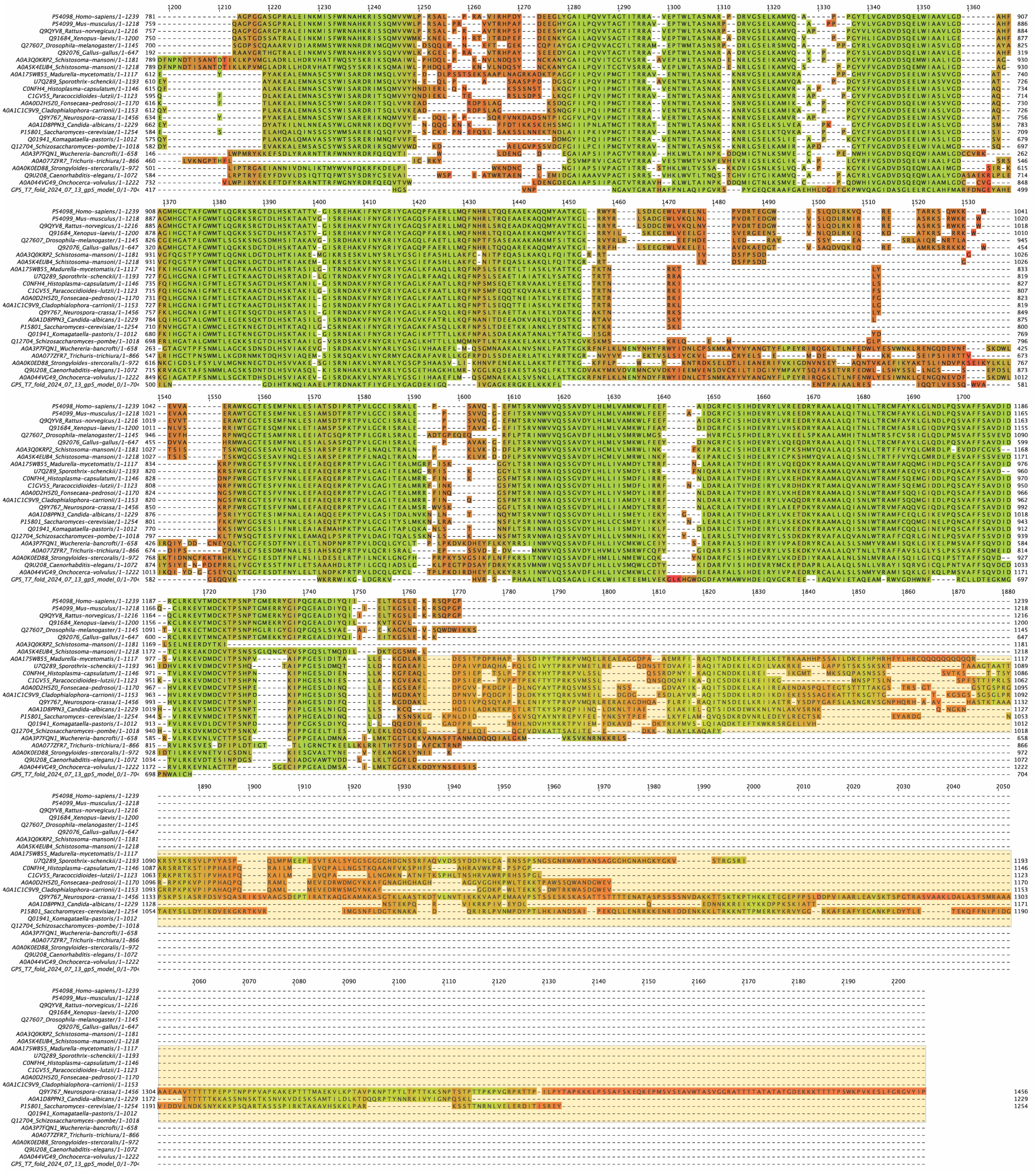

b

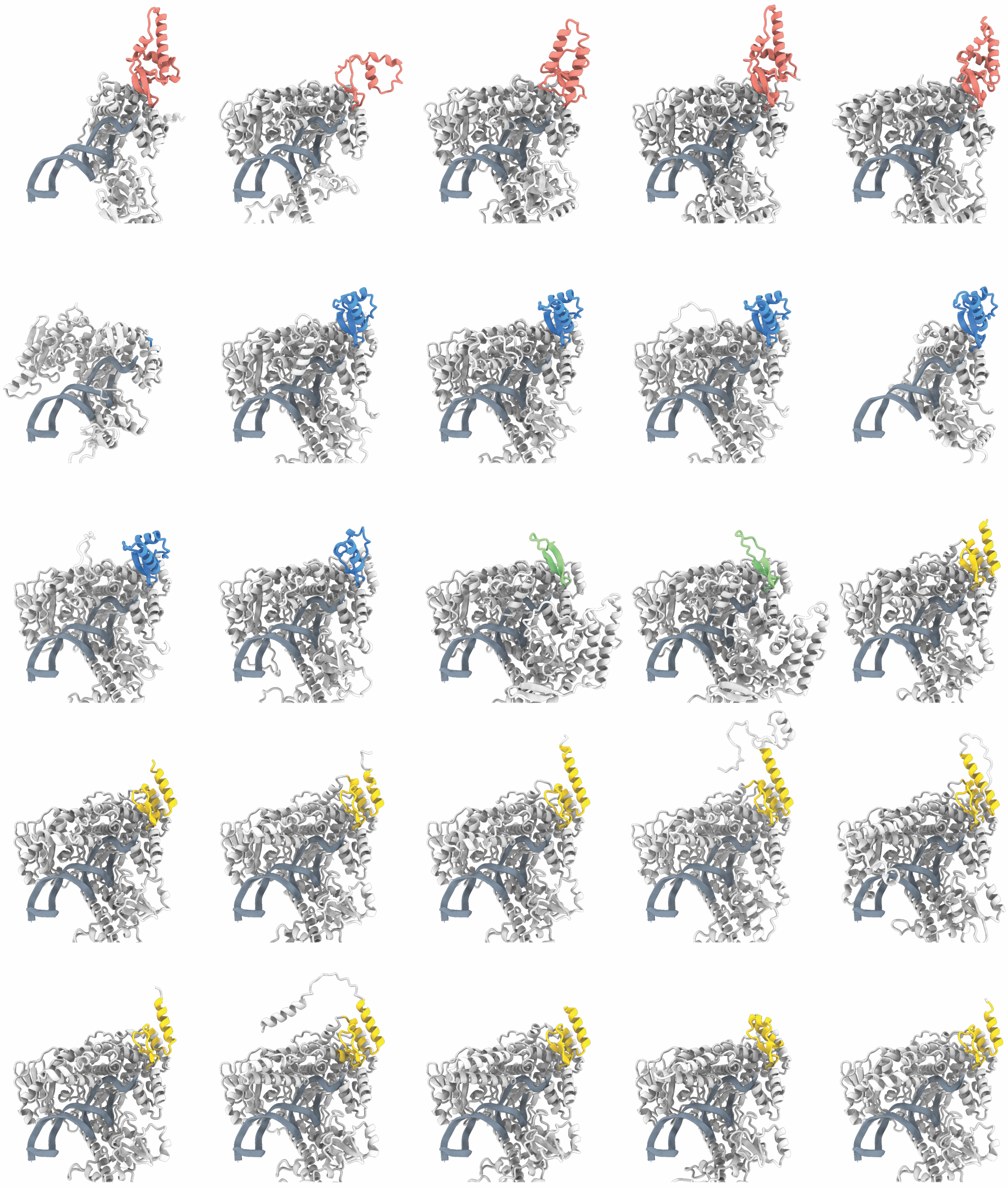

**Supplementary Figure 8. Sequence and structural diversity of catcher-like insertions across mitochondrial DNA polymerases. a,** Multiple-sequence alignment representation of structural alignment from FoldSeek. Only top hits are shown. Lineage-specific insertions are indicated by the alignment colouring. **b,** Structural representation of AlphaFold predictions from each sequence showed in the alignment. Structures are displayed in a common orientation, with the polymerase core shown in grey and lineage-specific insertions coloured as in Supplementary Figure 7.

### 2. SUPPLEMENTARY TABLES

**Supplementary Table 1: Cryo-EM data collection, refinement and validation statistics**

| **General Info** | | | | | | |
|---|---|---|---|---|---|---|
| DATASET ID | **Polγ-DNA-mtSSB** | **Polγ-G4a** | **Polγ-G4b** | **Polγ-SD-open** | **Polγ-SD-close** | **Polγ-APO** |
| EMDB | EMD-XXXXX | EMD-XXXXX | EMD-XXXXX | EMD-XXXXX | EMD-XXXXX | EMD-XXXXX |
| PDB | XXXX | XXXX | XXXX | XXXX | - | - |
| **Data collection and processing** | | | | | | |
| Magnification | 130000 | 130000 | 130000 | 105000 | 105000 | 105000 |
| Voltage (kV) | 300 | 300 | 300 | 300 | 300 | 300 |
| Electron exposure ($e^{-}$/Å$^2$) | 1 | 1,003 | 1,003 | 1,11 | 1,11 | 1,11 |
| Defocus range (μm) | 0.8-2.5 | 0,8-2,5 | 0,8-2,5 | 0.8-2.5 | 0.8-2.5 | 0.8-2.5 |
| Pixel size (Å) | 0,653 | 0,645 | 0,645 | 0,8238 | 0,8238 | 0,8238 |
| Symmetry imposed | C1 | C1 | C1 | C1 | C1 | C1 |
| Initial particle images (no.) | 1469483 | 1000941 | 1000941 | 3594509 | 3594509 | 3594509 |
| Final particle images (no.) | 170136 | 209158 | 68861 | 225443 | 836840 | 984666 |
| Map resolution (FSC=0.143) | 2,4 | 3,2 | 3,4 | 3,1 | 3,7 | 3,2 |
| **Refinement** | | | | | | |
| Initial model used | PDB 5C52 | Polγ-DNA-mtSSB | Polγ-G4a | Polγ-DNA-mtSSB | | |
| Model resolution (FSC=0.5, Å) | 2,6 | 6,6 | 7,7 | 4,1 | | |
| Model composition | | | | | | |
| Non-hydrogen atoms | 15230 | 15201 | 14602 | 14249 | | |
| Protein residues | 1791 | 1791 | 1739 | 1716 | | |
| Ligands (DNA) | 40 | 30 | 31 | 21 | | |
| Ligands | 1 (Mg, DDS) | 1 (Mg) | 1 (Mg) | 1 (DCP) | | |
| B factors (min-max, Å$^2$) | 40.00-227.71 | 101.52-1021.10 | 123.09-644.00 | 126.25-624.30 | | |
| R.m.s. deviations | | | | | | |
| Bond lengths (Å) | 0,4 | 0,00406 | 0,02613 | 0,0038 | | |
| Bond angles (°) | 0,64 | 0,85502 | 2,01906 | 0,84329 | | |
| **Validation** | | | | | | |
| MolProbity score | 1,5 | 2,85 | 2,81 | 1,63 | | |
| Clashscore | 4,61 | 73,93 | 48,52 | 7,74 | | |
| Poor rotamers (%) | 0,77 | 1,59 | 2,05 | 0,47 | | |
| Ramachandran plot | | | | | | |
| Favored (%) | 96,16 | 94,76 | 93,94 | 96,68 | | |
| Allowed (%) | 3,79 | 5,01 | 5,59 | 3,26 | | |
| Disallowed (%) | 0,06 | 0,23 | 0,47 | 0,06 | | |

**Supplementary Table 2: DNA Substrates**

| **cryoEM Substrates** | **Oligonucleotide** | **Sequence (5'-3')** |
|---|---|---|
| **60ssDNA** | 60ssDNA-1 | AGGCTCGCTcGcTCGCTCGGATTTTTTTTTTTTTTTTTTTTTTTTTTTTTTTTTTTTTTTTTTTTTTTTTTTTTTTTTTTTTTAGGCTCGCTcGcTCGCTCGGA |
| | 60ssDNA-2 | TCCGAGCGAgCgAGCGAGCCT |
| | 60ssDNA-3 | TCCGAGCGAgCgAGCGAGCCT |
| **SD-DNA-cryo** | SD-DNA-cryo-1 | CGCCAGGGTTTTCCCAGT*C*A*C |
| | SD-DNA-cryo-2 | GCACTGGCCGTCGTTTTACGAAGAGGGAGGTGACTGGGAAAACCCTG*G*C*G |
| | SD-DNA-cryo-3 | ACTTGAATGCGGCTTAGTATGCATTGTAAAACGACGGCCAG*T*G*C |
| **G4b-DNA** | G4b-DNA-1 | CGCCAGGGTTTTC*C*G |
| | G4b-DNA-2 | TAAACTGTGGGGGGTGTCTTTGGGGTTTGGTTGGTTCGGGGTATGGGGTTAGCAGCGCGGAAAACCCTGGCG |
| **Biochemistry Substrates** | **Oligonucleotide** | **Sequence (5'-3')** |
| **G4a (G4-5 in Sullivan et. al. (2020) J. Biol. Chem.)** | G4a-1 | **IRD800**-CGCCAGGGTTTTC*C*G |
| | G4a-2 | TAAACTGTGGGGGGTGTCTTTGGGGTTTGGTTGGTTCGGGGTATGGGGTTAGCAGCGCGGAAAACCCTGGCG |
| **G4b (G4-8 in Sullivan et. al. (2020) J. Biol. Chem.)** | G4b-1 | **IRD800**-CGCCAGGGTTTTC*C*G |
| | G4b-2 | CTAGAGGGGGTAGAGGGGGGTGCTATAGGGTAAATACGGGCCCTATTTCCGGAAAACCCTGGCG |
| **SD-RNA-flap** | SD-RNA-flap-1 | **Cy5**-CGCCAGGGTTTTCCCAGT*C*A*C |
| | SD-RNA-flap-2 | GCACTGGCCGTCGTTTTACGAAGAGGGAGGTGACTGGGAAAACCCTG*G*C*G |
| | SD-RNA-flap-3 | ACUUGAAUGCGGCUUAGUAUGCAUUGUAAAACGACGGCCAGU*G*C |
| **long-PE-DNA** | long-PE-DNA-1 | **IRD800**-CGCCAGGGTTTTC*C*G |
| | long-PE-DNA-2 | ACACGAACTGGATGCTACCTGAAGTGATTGATTACGATGAACGTGAACTGGATGCTACCTATTAGACTGAACACGAACTGGggtaccATGCTACCTGAAGTGATTGATTACGATGAACATGAACTCGATGCTACCTGAAGTGATTGGACTCGGAAAACCCTGGCG |
| **SD-DNA** | SD-DNA-1 | **IRD700**-CGCCAGGGTTTTCCCAGTCAC |
| | SD-DNA-2 | GCACTGGCCGTCGTTTTACGAAGAGGGAGGTGACTGGGAAAACCCTG*G*C*G |
| | SD-DNA-3 | ACTTGAATGCGGCTTAGTATGCATTGTAAAACGACGGCCAG*T*G*C |
| **PE-DNA** | PE-DNA-1 | **IRD700**-CGCCAGGGTTTTCCCAGTCAC |
| | PE-DNA-2 | GCACTGGCCGTCGTTTTACGAAGAGGGAGGTGACTGGGAAAACCCTG*G*C*G |
| **CE Substrates** | | **oligo 1 (5'-3')** |
| **SD-DNA-CE** | SD-DNA-CE-1 | **6-FAM**-CGCCAGGGTTTTCCCAGTCAC |
| | SD-DNA-CE-2 | GCACTGGCCGTCGTTTTACGAAGAGGGAGGTGACTGGGAAAACCCTG*G*C*G |
| | SD-DNA-CE-3 | ACTTGAATGCGGCTTAGTATGCATTGTAAAACGACGGCCAG*T*G*C |
| **control-DNA** | control-DNA-1 | **6-FAM**-ctgTTAGGGTTAGGGTTAAAATTAGGGTTAGGGTTAGtatcgtctGTTAGGGTTAG |
| **FP Substrates** | | **oligo 1 (5'-3')** |
| **Forked DNA** | Forked-DNA-1 | **6-FAM**-ACTTGAATGCGGCTTAGTATGCATTGTAAAACGACGGCCAGTGC |
| | Forked-DNA-2 | GCACTGGCCGTCGTTTTACGGTCGTGACTGGGAAAACCCTGGCG |
| **Primer-Template-DNA** | Primer-Template-DNA-1 | **6-FAM**-AATTATTTATCCTACGCGCCCCTCCTAGATGACACCGCCAGGGTTTTCC*G |
| | Primer-Template-DNA-2 | CTGAAGTGATTGGACTCGGAAAACCCTGGCGGTGTCATCTAGGAGGGGCGCGTAGGATAAATAATT |
| **ssDNA** | ssDNA-1 | **6-FAM**-ACTTGAATGCGGCTTAGTATGCATTGTAAAACGACGGCCAGTGC |

**Supplementary Table 3: DNA Constructs**

| Protein | Variant | Construct |
|---|---|---|
| PolγA | WT | pACEBac1-PolG1(30-1239)-3xFlag |
| mtSSB | WT | pRSFDuetT5-10xHis-SSB(17-148)-StrepTag |
| PolγB | WT | pETite-6xHis-SUMO-PolG2(26-485) |
| PolγA | R1006A | pACEBac1-PolG1(30-1239)-3xFlag-R1006A |
| PolγA | E1016A | pACEBac1-PolG1(30-1239)-3xFlag-E1016A |
| PolγA | R1026A | pACEBac1-PolG1(30-1239)-3xFlag-R1026A |
| PolγA | R1026A-R1030A-R1034A (RRR) | pACEBac1-PolG1(30-1239)-3xFlag-R1026A-R1030A-R1034A |
| PolγA | R1030A | pACEBac1-PolG1(30-1239)-3xFlag-R1030A |
| PolγA | R1034A | pACEBac1-PolG1(30-1239)-3xFlag-R1034A |
| PolγA | W1038A | pACEBac1-PolG1(30-1239)-3xFlag-W1038A |
| PolγA | W1041A | pACEBac1-PolG1(30-1239)-3xFlag-W1041A |
| PolγA | L992 to G1051 (Δ991-1051) | pACEBac1-PolG1(30-1239)-3xFlag-dDL |
| PolγA | D274A | pACEBac1-PolG1(30-1239)-3xFlag -D274A |
| PolγA | 991-1051 WT | pRSFDuetT5_H10-MBP-TEV-PolGa(991-1051) |
| PolγA | 991-1051 R1026A | pRSFDuetT5_H10-MBP-TEV-PolG1(991-1051)-3C-Strep-R1026A |
| PolγA | 991-1051 R1030A | pRSFDuetT5_H10-MBP-TEV-PolG1(991-1051)-3C-Strep-R1030A |
| PolγA | 991-1051 R1047A | pRSFDuetT5_H10-MBP-TEV-PolG1(991-1051)-3C-Strep-R1047A |
| PolγA | 991-1051 R1026A-R1030A-R1034A (RRR) | pRSFDuetT5_H10-MBP-TEV-PolG1(991-1051)-R1026A-R1030A-R1034A |

**Supplementary Table 4. FoldSeek results**

| Target | Database | Description | Species | % ID | E-value | Score | Query Position | Target Position |
|---|---|---|---|---|---|---|---|---|
| F8W5R6 | AFDB-PROTEOME | DNA polymerase subunit gamma-1 | Danio-rerio | 58.1 | 0.00E+00 | 4940 | 2-865 (869) | 37-1197 (1206) |
| P54098 | AFDB-PROTEOME | DNA polymerase subunit gamma-1 | Homo-sapiens | 73.8 | 0.00E+00 | 5605 | 1-865 (869) | 67-1233 (1239) |
| P54099 | AFDB-PROTEOME | DNA polymerase subunit gamma-1 | Mus-musculus | 69.1 | 0.00E+00 | 5397 | 1-865 (869) | 50-1212 (1218) |
| Q9QYV8 | AFDB-PROTEOME | DNA polymerase subunit gamma-1 | Rattus-norvegicus | 68.4 | 0.00E+00 | 5370 | 1-865 (869) | 50-1210 (1216) |
| Q91684 | AFDB-SWISSPROT | DNA polymerase subunit gamma-1 | Xenopus-laevis | 57.2 | 0.00E+00 | 4823 | 1-862 (869) | 41-1199 (1200) |
| Q27607 | AFDB-PROTEOME | DNA polymerase subunit gamma-1, mitochondrial | Drosophila-melanogaster | 41.8 | 2.70E-87 | 3708 | 2-857 (869) | 50-1129 (1145) |
| A0A175WB55 | AFDB-PROTEOME | Mitochondrial DNA polymerase catalytic subunit | Madurella-mycetomatis | 41.1 | 3.56E-80 | 3358 | 3-865 (869) | 58-1015 (1117) |
| C1GV55 | AFDB-PROTEOME | Mitochondrial DNA polymerase catalytic subunit | Paracoccidioides-lutzii-Pb01 | 40 | 8.11E-79 | 3311 | 3-865 (869) | 42-989 (1123) |
| Q9Y767 | AFDB-SWISSPROT | DNA polymerase gamma, mitochondrial | Neurospora-crassa-OR74A | 39.6 | 1.61E-78 | 3285 | 3-865 (869) | 59-1031 (1456) |
| U7Q289 | AFDB-PROTEOME | Mitochondrial DNA polymerase catalytic subunit | Sporothrix-schenckii-ATCC-58251 | 40.5 | 2.19E-78 | 3327 | 2-868 (869) | 56-1012 (1193) |
| A0A1C1C9V9 | AFDB-PROTEOME | Mitochondrial DNA polymerase catalytic subunit | Cladophialophora-carrionii | 41.3 | 1.94E-77 | 3230 | 3-864 (869) | 59-1003 (1153) |
| C0NFH4 | AFDB-PROTEOME | Mitochondrial DNA polymerase catalytic subunit | Histoplasma-capsulatum-G186AR | 39.9 | 2.13E-77 | 3259 | 3-864 (869) | 62-1011 (1146) |
| A0A5K4EUB4 | AFDB-PROTEOME | Mitochondrial DNA polymerase catalytic subunit | Schistosoma-mansoni | 32.7 | 4.14E-77 | 3271 | 1-865 (869) | 30-1218 (1218) |
| A0A0D2H5Z0 | AFDB-PROTEOME | Mitochondrial DNA polymerase catalytic subunit | Fonsecaea-pedrosoi-CBS-271.37 | 41.3 | 6.05E-77 | 3207 | 3-865 (869) | 63-1005 (1170) |
| Q12704 | AFDB-PROTEOME | DNA polymerase gamma | Schizosaccharomyces-pombe-972h- | 38.6 | 6.77E-76 | 3215 | 1-864 (869) | 40-979 (1018) |
| Q01941 | AFDB-SWISSPROT | DNA polymerase gamma | Komagataella-pastoris | 37.5 | 6.72E-75 | 3106 | 3-865 (869) | 17-953 (1012) |
| A0A1D8PPN3 | AFDB-PROTEOME | Mitochondrial DNA polymerase catalytic subunit | Candida-albicans-SC5314 | 36.4 | 3.14E-74 | 3074 | 1-869 (869) | 49-1065 (1229) |
| P15801 | AFDB-PROTEOME | DNA polymerase gamma | Saccharomyces-cerevisiae-S288C | 37.9 | 1.81E-73 | 3039 | 3-865 (869) | 39-984 (1254) |
| A0A3Q0KRP2 | AFDB-PROTEOME | Mitochondrial DNA polymerase catalytic subunit | Schistosoma-mansoni | 32 | 1.30E-68 | 2806 | 1-824 (869) | 30-1175 (1181) |
| A0A044VG49 | AFDB-PROTEOME | Mitochondrial DNA polymerase catalytic subunit | Onchocerca-volvulus | 31.9 | 2.48E-64 | 2580 | 10-865 (869) | 199-1211 (1222) |
| A0A158Q5M0 | AFDB-PROTEOME | Mitochondrial DNA polymerase catalytic subunit | Dracunculus-medinensis | 31.8 | 9.97E-63 | 2478 | 10-852 (869) | 201-1205 (1213) |
| A0A0K0ED88 | AFDB-PROTEOME | Mitochondrial DNA polymerase catalytic subunit | Strongyloides-stercoralis | 30.6 | 2.96E-62 | 2452 | 54-865 (869) | 1-969 (972) |
| Q9U208 | AFDB-PROTEOME | Mitochondrial DNA polymerase catalytic subunit | Caenorhabditis-elegans | 29.9 | 2.01E-60 | 2367 | 1-864 (869) | 23-1072 (1072) |
| A0A077ZFR7 | AFDB-PROTEOME | Mitochondrial DNA polymerase catalytic subunit | Trichuris-trichiura | 37 | 2.55E-60 | 2162 | 32-858 (869) | 2-848 (866) |
| Q92076 | AFDB-SWISSPROT | DNA polymerase subunit gamma-1 | Gallus-gallus | 60.8 | 3.59E-53 | 1844 | 405-865 (869) | 1-646 (647) |
| A0A3P7FQN1 | AFDB-PROTEOME | Mitochondrial DNA polymerase catalytic subunit | Wuchereria-bancrofti | 31.9 | 3.64E-31 | 1008 | 415-865 (869) | 20-624 (658) |
| A0A3P7E5A7 | AFDB-PROTEOME | DNA-directed DNA polymerase | Wuchereria-bancrofti | 33.1 | 3.57E-22 | 773 | 12-375 (869) | 48-405 (410) |
| Q9HT80 | AFDB-PROTEOME | DNA polymerase I | Pseudomonas-aeruginosa-PAO1 | 14.8 | 3.77E-14 | 319 | 56-820 (869) | 253-908 (913) |
| A0A077ZEH0 | AFDB-PROTEOME | DNA-directed DNA polymerase | Trichuris-trichiura | 14.1 | 8.44E-14 | 331 | 125-795 (869) | 348-907 (928) |
| K7M4K7 | AFDB-PROTEOME | POLAc domain-containing protein | Glycine-max | 14.7 | 9.28E-14 | 286 | 48-822 (869) | 245-1068 (1074) |
| A0A0H3GPI6 | AFDB-PROTEOME | DNA polymerase I | Klebsiella-pneumoniae-subsp.-pneumoniae-HS11286 | 14.2 | 9.73E-14 | 290 | 33-795 (869) | 239-871 (892) |
| P00582 | AFDB-PROTEOME | DNA polymerase I | Escherichia-coli-K-12 | 14.2 | 1.23E-13 | 319 | 125-795 (869) | 348-907 (928) |
| Q32A35 | AFDB-PROTEOME | DNA polymerase I | Shigella-dysenteriae-Sd197 | 13.8 | 1.42E-13 | 327 | 125-795 (869) | 348-907 (928) |
| P56105 | AFDB-PROTEOME | DNA polymerase I | Helicobacter-pylori-26695 | 14.1 | 1.72E-13 | 304 | 28-820 (869) | 265-885 (891) |
| Q0PBH1 | AFDB-PROTEOME | DNA polymerase I | Campylobacter-jejuni-subsp.-jejuni-NCTC-11168-=-ATCC-700819 | 14.3 | 1.89E-13 | 323 | 123-820 (869) | 313-874 (879) |
| Q9F173 | AFDB-PROTEOME | DNA polymerase I | Salmonella-enterica-subsp.-enterica-serovar-Typhimurium-str.-LT2 | 13.8 | 1.89E-13 | 283 | 41-795 (869) | 290-907 (928) |
| Q6Z4T3 | AFDB-PROTEOME | DNA polymerase I B, mitochondrial | Oryza-sativa-Japonica-Group | 14.6 | 2.28E-13 | 274 | 49-820 (869) | 210-1030 (1035) |
| Q5F533 | AFDB-PROTEOME | DNA polymerase I | Neisseria-gonorrhoeae-FA-1090 | 14.7 | 3.34E-13 | 315 | 125-820 (869) | 350-925 (930) |
| Q7TQ07 | AFDB-PROTEOME | DNA polymerase nu | Mus-musculus | 13.3 | 3.84E-13 | 260 | 1-820 (869) | 103-850 (864) |
| Q84ND9 | AFDB-PROTEOME | DNA polymerase I B, chloroplastic/mitochondrial | Arabidopsis-thaliana | 15 | 4.43E-13 | 269 | 48-822 (869) | 182-1028 (1034) |
| A0A1D6IZ69 | AFDB-PROTEOME | DNA polymerase like1 | Zea-mays | 14.1 | 5.11E-13 | 271 | 48-822 (869) | 227-1033 (1039) |
| F1RDP8 | AFDB-PROTEOME | Polymerase (DNA directed) nu | Danio-rerio | 12.3 | 6.48E-13 | 256 | 48-820 (869) | 454-1128 (1146) |
| I1JJ76 | AFDB-PROTEOME | POLAc domain-containing protein | Glycine-max | 14.2 | 6.48E-13 | 265 | 48-822 (869) | 259-1071 (1077) |
| Q4DT24 | AFDB-PROTEOME | DNA-directed DNA polymerase | Trypanosoma-cruzi-strain-CL-Brener | 13.8 | 6.48E-13 | 251 | 24-820 (869) | 226-908 (924) |
| F1LWX6 | AFDB-PROTEOME | DNA polymerase nu | Rattus-norvegicus | 13.5 | 6.79E-13 | 250 | 1-820 (869) | 101-848 (862) |
| Q583X2 | AFDB-PROTEOME | DNA-directed DNA polymerase | Trypanosoma-brucei-brucei-TREU927 | 13.4 | 6.79E-13 | 280 | 124-820 (869) | 271-939 (958) |
| A4I9T9 | AFDB-PROTEOME | DNA-directed DNA polymerase | Leishmania-infantum | 12.8 | 1.04E-12 | 259 | 143-823 (869) | 505-1244 (1257) |
| P43741 | AFDB-PROTEOME | DNA polymerase I | Haemophilus-influenzae-Rd-KW20 | 14.5 | 1.32E-12 | 304 | 126-792 (869) | 353-913 (930) |

| | | | | | | | | |
|---|---|---|---|---|---|---|---|---|
| A0A1D6JFD6 | AFDB-PROTEOME | White seedling2 | Zea-mays | 15.3 | 1.32E-12 | 247 | 48-822 (869) | 212-1046 (1052) |
| Q385L3 | AFDB-PROTEOME | DNA-directed DNA polymerase | Trypanosoma-brucei-brucei-TREU927 | 12.8 | 2.02E-12 | 243 | 128-820 (869) | 176-886 (896) |
| F4I6M1 | AFDB-PROTEOME | DNA polymerase I A, chloroplastic/mitochondrial | Arabidopsis-thaliana | 14.4 | 2.12E-12 | 265 | 48-823 (869) | 209-1045 (1050) |
| Q6Z4T5 | AFDB-PROTEOME | DNA polymerase I A, chloroplastic | Oryza-sativa-Japonica-Group | 14.6 | 2.22E-12 | 257 | 49-817 (869) | 214-1023 (1033) |
| Q7Z5Q5 | AFDB-PROTEOME | DNA polymerase nu | Homo-sapiens | 12.1 | 3.24E-12 | 253 | 48-820 (869) | 148-851 (900) |
| O18475 | AFDB-PROTEOME | DNA polymerase theta | Drosophila-melanogaster | 11.8 | 3.74E-12 | 246 | 25-820 (869) | 1326-2050 (2059) |
| K0EPY7 | AFDB-PROTEOME | Bifunctional 3'-5' exonuclease/DNA polymerase | Nocardia-brasiliensis-ATCC-700358 | 14.3 | 4.11E-12 | 288 | 128-820 (869) | 2-550 (555) |
| Q4E329 | AFDB-PROTEOME | DNA-directed DNA polymerase | Trypanosoma-cruzi-strain-CL-Brener | 13.1 | 5.72E-12 | 247 | 24-820 (869) | 229-911 (927) |
| Q4DV47 | AFDB-PROTEOME | DNA polymerase theta (Polymerase domain), putative | Trypanosoma-cruzi-strain-CL-Brener | 12.8 | 7.25E-12 | 251 | 215-820 (869) | 279-872 (881) |
| D4A628 | AFDB-PROTEOME | DNA-directed DNA polymerase | Rattus-norvegicus | 13.2 | 9.19E-12 | 240 | 48-820 (869) | 1768-2538 (2547) |
| F1Q4P4 | AFDB-PROTEOME | Polymerase (DNA directed), theta | Danio-rerio | 12.2 | 9.64E-12 | 228 | 48-820 (869) | 1800-2567 (2576) |
| A0A5P3FTI1 | AFDB-PROTEOME | DNA polymerase I | Enterococcus-faecium | 14.3 | 1.11E-11 | 237 | 66-820 (869) | 269-876 (881) |
| P9WNU5 | AFDB-PROTEOME | DNA polymerase I | Mycobacterium-tuberculosis-H37Rv | 14.2 | 1.48E-11 | 265 | 125-820 (869) | 336-898 (904) |
| K0ESK5 | AFDB-PROTEOME | DNA polymerase I | Nocardia-brasiliensis-ATCC-700358 | 12.4 | 1.48E-11 | 266 | 128-820 (869) | 321-881 (887) |
| P46835 | AFDB-PROTEOME | DNA polymerase I | Mycobacterium-leprae-TN | 13.1 | 1.70E-11 | 277 | 124-820 (869) | 338-905 (911) |
| Q8CGS6 | AFDB-PROTEOME | DNA polymerase theta | Mus-musculus | 11.8 | 2.49E-11 | 241 | 126-820 (869) | 1802-2535 (2544) |
| Q8I7W9 | AFDB-PROTEOME | DNA-directed DNA polymerase | Dictyostelium-discoideum | 13 | 4.60E-11 | 236 | 32-818 (869) | 566-1360 (1369) |
| P59200 | AFDB-PROTEOME | DNA polymerase I | Streptococcus-pneumoniae-R6 | 12.9 | 5.06E-11 | 248 | 66-804 (869) | 268-861 (877) |
| A0A044UT20 | AFDB-PROTEOME | DNA-directed DNA polymerase | Onchocerca-volvulus | 11.5 | 7.05E-11 | 259 | 125-820 (869) | 1195-1750 (1760) |
| Q2FXN9 | AFDB-PROTEOME | DNA polymerase I | Staphylococcus-aureus-subsp.-aureus-NCTC-8325 | 12.1 | 7.75E-11 | 249 | 70-820 (869) | 271-871 (876) |
| A0FLQ6 | AFDB-PROTEOME | DNA polymerase theta | Caenorhabditis-elegans | 14.1 | 1.08E-10 | 260 | 246-820 (869) | 1170-1656 (1661) |
| Q8ILY1 | AFDB-PROTEOME | Plastid replication-repair enzyme | Plasmodium-falciparum-3D7 | 10.4 | 1.08E-10 | 267 | 124-820 (869) | 1462-2011 (2016) |
| O75417 | AFDB-PROTEOME | DNA polymerase theta | Homo-sapiens | 12.5 | 1.58E-10 | 198 | 49-825 (869) | 1803-2590 (2590) |
| A0A1D6IVX5 | AFDB-PROTEOME | Helicase and polymerase-containing protein TEBICHI | Zea-mays | 11.1 | 4.92E-10 | 227 | 49-820 (869) | 1362-2082 (2094) |
| Q54S42 | AFDB-PROTEOME | POLAc domain-containing protein | Dictyostelium-discoideum | 12.5 | 5.41E-10 | 226 | 188-820 (869) | 786-1426 (1459) |
| X8FFS9 | AFDB-PROTEOME | DNA polymerase I | Mycobacterium-ulcerans-str.-Harvey | 15.8 | 8.28E-10 | 255 | 347-820 (869) | 7-399 (405) |
| K7M6P7 | AFDB-PROTEOME | Uncharacterized protein | Glycine-max | 12.4 | 9.55E-10 | 239 | 125-820 (869) | 1457-2126 (2147) |
| A0A0N4U9W9 | AFDB-PROTEOME | DNA-directed DNA polymerase | Dracunculus-medinensis | 10.2 | 1.00E-09 | 226 | 125-823 (869) | 352-932 (940) |
| A0A3Q0KQI9 | AFDB-PROTEOME | DNA-directed DNA polymerase | Schistosoma-mansoni | 11.9 | 1.00E-09 | 195 | 47-820 (869) | 1401-2216 (2225) |
| Q0J9P9 | AFDB-PROTEOME | Os04g0637400 protein | Oryza-sativa-Japonica-Group | 16.6 | 1.05E-09 | 227 | 315-823 (869) | 28-555 (560) |
| Q86KS9 | AFDB-PROTEOME | DNA-directed DNA polymerase | Dictyostelium-discoideum | 13 | 1.15E-09 | 243 | 287-820 (869) | 2-492 (516) |
| Q4D4N5 | AFDB-PROTEOME | DNA-directed DNA polymerase | Trypanosoma-cruzi-strain-CL-Brener | 10.3 | 1.94E-09 | 199 | 35-820 (869) | 453-1428 (1473) |
| Q588V7 | AFDB-PROTEOME | Helicase and polymerase-containing protein TEBICHI | Arabidopsis-thaliana | 11.3 | 6.98E-09 | 171 | 48-820 (869) | 1406-2144 (2154) |
| Q4DSW2 | AFDB-PROTEOME | DNA-directed DNA polymerase | Trypanosoma-cruzi-strain-CL-Brener | 11.2 | 7.68E-09 | 184 | 42-820 (869) | 400-1372 (1417) |
| A0A0P0Y964 | AFDB-PROTEOME | Os12g0291000 protein | Oryza-sativa-Japonica-Group | 12.5 | 4.43E-08 | 194 | 126-795 (869) | 1403-2043 (2051) |
| Q2FYA1 | AFDB-PROTEOME | DNA-directed DNA polymerase | Staphylococcus-aureus-subsp.-aureus-NCTC-8325 | 9.9 | 5.11E-08 | 214 | 128-824 (869) | 2-650 (650) |
| E9AGL3 | AFDB-PROTEOME | Mitochondrial_DNA_polymerase_I_protein_C_-_putative | Leishmania-infantum | 10.2 | 3.24E-07 | 153 | 15-820 (869) | 449-1542 (1548) |
| Q4D7R3 | AFDB-PROTEOME | Mitochondrial DNA polymerase I protein C, putative | Trypanosoma-cruzi-strain-CL-Brener | 10.3 | 4.11E-07 | 141 | 24-820 (869) | 576-1621 (1627) |
| Q86HT8 | AFDB-PROTEOME | POLAc domain-containing protein | Dictyostelium-discoideum | 11 | 1.11E-06 | 179 | 333-829 (869) | 3-497 (519) |
| C6KT89 | AFDB-PROTEOME | DNA polymerase 1, putative | Plasmodium-falciparum-3D7 | 9.7 | 1.34E-06 | 164 | 386-800 (869) | 931-1354 (1444) |
| Q57UM9 | AFDB-PROTEOME | DNA-directed DNA polymerase | Trypanosoma-brucei-brucei-TREU927 | 13.4 | 1.78E-06 | 182 | 451-820 (869) | 1271-1643 (1649) |
| K0ERU8 | AFDB-PROTEOME | Diphosphomevalonate decarboxylase | Nocardia-brasiliensis-ATCC-700358 | 12.9 | 3.88E+00 | 26 | 714-785 (869) | 221-313 (339) |
| K0F4I6 | AFDB-PROTEOME | Diphosphomevalonate decarboxylase | Nocardia-brasiliensis-ATCC-700358 | 10.7 | 4.07E+00 | 25 | 718-785 (869) | 226-318 (350) |
